# Defining the lineage of thermogenic perivascular adipose tissue

**DOI:** 10.1101/2020.06.23.164772

**Authors:** Anthony R. Angueira, Alexander P. Sakers, Lan Cheng, Rojesh Shrestha, Chihiro Okada, Kirill Batmanov, Katalin Susztak, Patrick Seale

## Abstract

Brown adipose tissue can expend large amounts of energy and thus increasing its amount or activity is a promising therapeutic approach to combat metabolic disease. In humans, major deposits of brown fat cells are found intimately associated with large blood vessels, corresponding to perivascular adipose tissue (PVAT). However, the cellular origins of PVAT are poorly understood. Here, we applied single cell transcriptomic analyses, *ex vivo* adipogenesis assays, and genetic fate mapping *in vivo* to determine the identity of perivascular adipocyte progenitors. We found that thoracic PVAT initially develops from a fibroblastic lineage, comprising progenitor cells (*Pdgfra*+;*Ly6a*+;*Pparg*−) and preadipocytes (*Pdgfra*+;*Ly6a*−;*Pparg*+). Progenitors and preadipocytes in PVAT shared transcriptional similarity with analogous cell types in white adipose tissue, pointing towards a conserved adipose cell lineage hierarchy. Interestingly, the aortic adventitia of adult animals contained a novel population of adipogenic smooth muscle cells (*Myh11*+; *Pdgfra*−; *Pparg*+) possessing the capacity to generate adipocytes *in vitro* and *in vivo*. Taken together, these studies define distinct populations of fibroblastic and smooth muscle progenitor cells for thermogenic PVAT, providing a crucial foundation for developing strategies to augment brown fat activity.

## Introduction

Thermogenic adipose tissue protects animals against hypothermia during cold exposure and is associated with resistance to obesity and metabolic syndrome in humans (Cypess et al., 2015; Harms and Seale, 2013; O’Mara et al., 2020; Yoneshiro et al., 2013). Thermogenic adipocytes have large numbers of mitochondria that contain Uncoupling Protein 1 (UCP1). Upon activation, UCP1 uncouples the mitochondrial proton gradient from ATP synthesis, creating an electrochemical driving force to burn large amounts of fatty acids and glucose for heat production (Cannon and Nedergaard, 2004). Additional thermogenic mechanisms, including creatine and calcium-driven futile cycles have also been described in adipocytes (Ikeda et al., 2017; Kazak et al., 2015, 2017). The therapeutic potential of thermogenic adipose tissue to suppress metabolic disease has spurred great interest in identifying the cellular origin and developmental pathways of thermogenic adipocytes.

Multiple discrete thermogenic adipose depots can be activated by cold exposure or β-adrenergic activators in humans (Cypess et al., 2009; van Marken Lichtenbelt et al., 2009; Sacks and Symonds, 2013; Virtanen et al., 2009). Infants have a thermogenic adipose tissue depot in the interscapular region, analogous to the location of the largest brown adipose tissue (BAT) depot in many rodent species (AHERNE and HULL, 1964; Harms and Seale, 2013; Heaton, 1972; Orava et al., 2011; Wang and Seale, 2016). However, the interscapular BAT pad regresses and becomes undetectable in adults (Lidell et al., 2013). Adult humans retain BAT deposits in the supraclavicular region and in several depots surrounding large blood vessels, known as perivascular adipose tissue (PVAT) (Cypess et al., 2013, 2015; Huttunen et al., 1981). For example, UCP1+ adipocytes are present within the carotid sheath and mediastinal cavity of humans (Cheung et al., 2013; Cypess et al., 2013). PVAT in the thoracic peri-aortic region of human subjects increase glucose uptake in response to systemic administration of a β3-adrenergic receptor agonist, a key functional attribute of BAT (Cypess et al., 2015). Therefore, understanding the ontogeny and regulation of thermogenic PVAT may identify novel therapeutic targets to combat metabolic disease.

Here, we performed a detailed analysis of peri-aortic adipose tissue development and maintenance in mice. We found that the aortic adipose depot forms perinatally and possesses characteristics of classical BAT, including: (1) stable expression of thermogenic components *in vivo* and *in vitro*; (2) dependence on the brown adipocyte-lineage determining factor EBF2; (3) responsiveness to environmental cold; and (4) resistance to high fat diet-induced inflammation. Single cell transcriptomic analysis of the aorta and associated adventitia in the perinatal period identified several distinct cell types, including presumptive preadipocytes, mesenchymal progenitor cells and three groups of SMCs. Cell differentiation assays and genetic lineage tracing studies show that fibroblastic progenitor cells but not SMCs are responsible for aortic PVAT organogenesis. Interestingly, the aortic adventitia of adult animals lacks fibroblastic preadipocytes but contains a novel population of adipogenic SMCs that can contribute to adipocyte formation. Together, these studies identify and define multiple progenitor cell types for thermogenic adipocytes in developing and adult PVAT.

## Results

### Aortic PVAT expresses an EBF2-dependent classical brown fat program

The aorta of adult mice is surrounded by multiple lobes of PVAT, all enclosed within an anatomic compartment defining fascial layer (Fig. 1A). Histologically, perivascular adipocytes have densely eosin-stained cytoplasm and contain multiple lipid droplets, resembling brown adipocytes in interscapular BAT (iBAT). PVAT from the thoracic aorta and iBAT expressed comparable levels of both common adipocyte genes (i.e. *Adipoq and Pparg2)* and the brown fat marker gene *Ucp1* (Fig. 1B). The thermogenic character of aortic PVAT was further recruited by cold exposure. Specifically, aortic PVAT from cold-exposed (4°C for 1 week) mice exhibited depleted lipid stores and expressed higher levels of thermogenesis-related genes (*Cycs, Ppargc1a,* and *Ucp1*), as compared to that from mice housed at thermoneutrality (30°C) (Fig. 1C,D).

**Fig.1.**
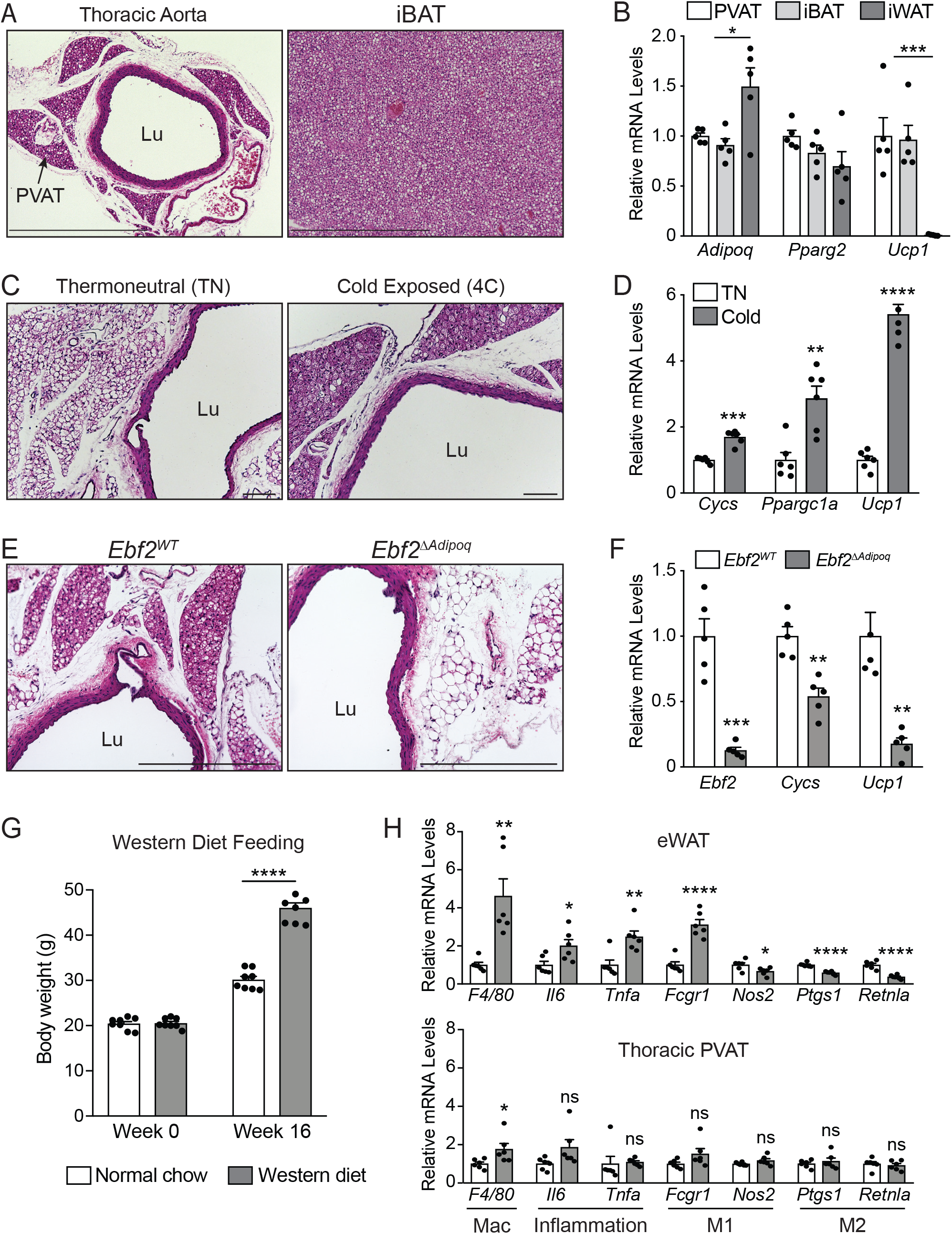
Aortic PVAT expresses an EBF2-regulated classical brown fat program. **(A)** H&E staining of thoracic aorta and iBAT from *C57/Bl6* mice (scale bar, 724.7 μm). **(B)** mRNA levels of indicated genes in thoracic PVAT, iBAT, and iWAT from *C57/Bl6* mice. (n=5 mice per group; mean+/− SEM). **(C, D)** H&E staining (C) and mRNA levels of indicated genes (D) in thoracic aorta from thermoneutrality-acclimated *C57/Bl6* mice housed at thermoneutrality or 4°C for one week. (n=6 mice per group; mean+/− SEM; scale bar, 100 μm). **(E, F)** H&E staining of thoracic aorta (E) and mRNA levels of indicated genes in aortic PVAT (F) from *Ebf2*^*WT*^ and *Ebf2*^*ΔAdipoq*^ mice (n=5 mice per group; mean+/− SEM; scale bar, 362.3 μm). **(G)** Body weights of *C57/Bl6* mice fed normal chow or western diet for 16 weeks (n=8 mice per group; mean+/−SEM) **(H)** mRNA levels of indicated genes in eWAT and thoracic PVAT from *C57/Bl6* mice fed normal chow or western diet for 16 weeks. (n=6 mice per group; mean+/− SEM). PVAT; Perivascular Adipose Tissue; Lu; vessel lumen

The transcription factor Early B Cell Factor-2 (EBF2) is a critical regulator of thermogenic adipocyte development (Rajakumari et al., 2013; Shapira et al., 2017; Stine et al., 2016; Angueira et al., 2020). To determine if EBF2 controls PVAT fate, we analyzed thoracic aortas from mice lacking *Ebf2* expression in adipocytes (*Ebf2*^*ΔAdipoq*^) and littermate control mice (*Ebf2*^*WT*^). Aortic PVAT from *Ebf2*^*ΔAdipoq*^ mice displayed a whitened morphology, associated with increased lipid deposition (Fig. 1E). *Ebf2* mutant tissue also showed markedly reduced expression of brown fat-specific genes, including an ~80% reduction of *Ucp1* and lower levels of *Cycs* (Fig. 1F). In contrast to WAT depots, iBAT is relatively resistant to inflammation triggered by a high fat or western diet (Dowal et al., 2017; Lumeng et al., 2007; Tian et al., 2016; Weisberg et al., 2003). To determine if PVAT is similarly protected from diet-induced inflammation, we fed *C57/Bl6* mice a western diet for 16 weeks (Fig 1G). As expected, western diet feeding promoted inflammation in epididymal WAT, typified by the higher expression levels of macrophage and pro-inflammatory genes (“M1-like macrophage”) and decreased levels of anti-inflammatory (“M2-like macrophage”) genes (Fig1 H). By contrast, western diet feeding did not affect the expression levels of various inflammatory or macrophage marker genes in thoracic PVAT. Altogether, these findings demonstrate that aortic PVAT possesses many of the phenotypic properties of classical BAT.

### Identification of fibroblast heterogeneity in developing aortic PVAT

We next sought to identify the progenitor cells responsible for the genesis of PVAT. Histologic examination of thoracic aorta from embryonic day 18 (E18) embryos (just prior to birth) revealed an extensive network of adventitial fibroblasts and well-developed aortic SMCs but an absence of lipid-containing adipocytes (Fig. 2A). During the perinatal period after birth (P1-P3), we found prominent lobes of multilocular adipocytes, organized in a stereotyped pattern around the aorta (Fig. 2A). Consistent with this histological (or anatomical) observation, there was a dramatic rise in the expression levels of adipocyte- and brown fat-marker genes (*Adipoq*, *Leptin* (*Lep*), *Pparg2*, *Ucp1*) in thoracic aortas between E18 and P3 (Fig. 2B).

**Fig. 2.**
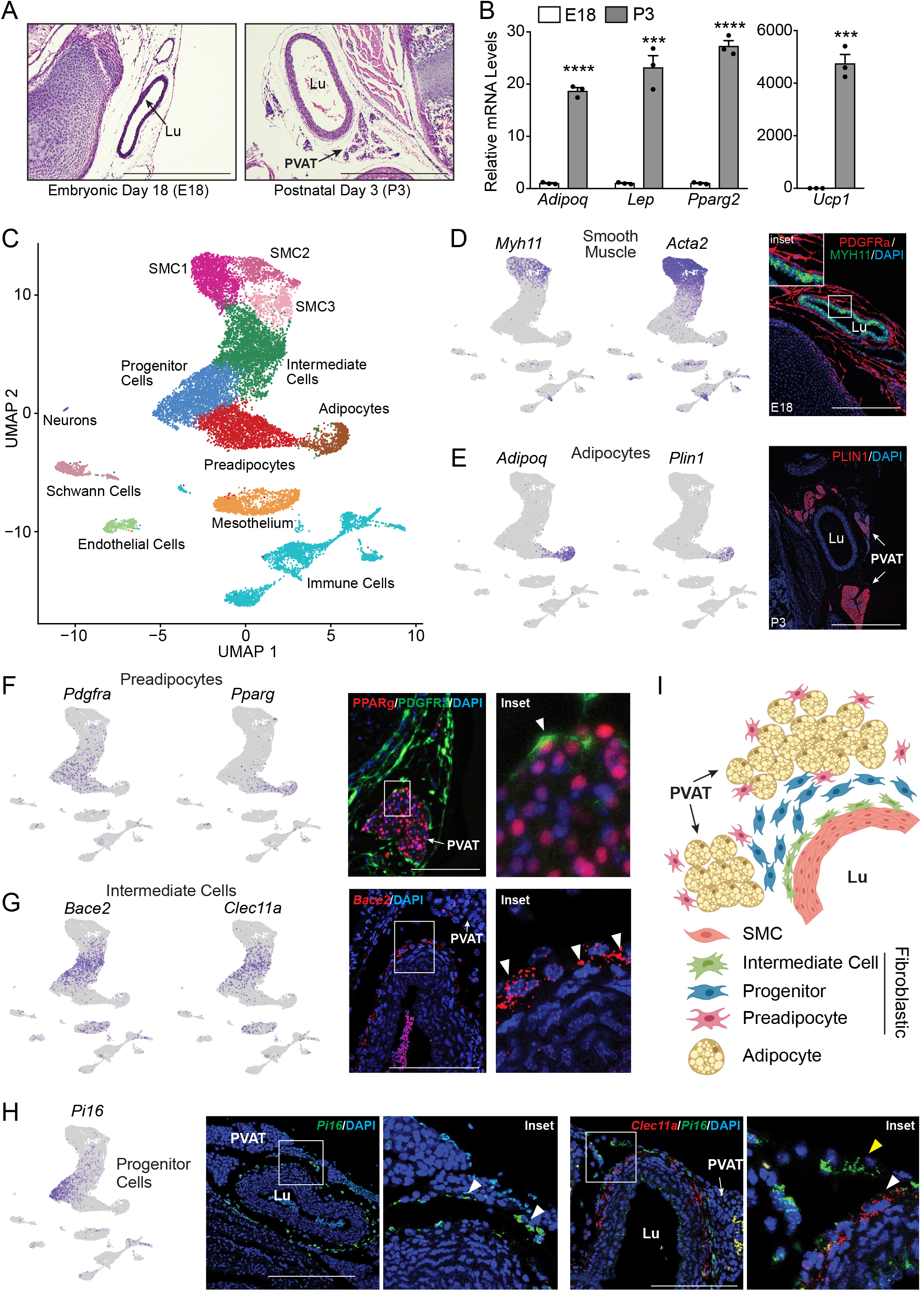
Identification of multiple fibroblast populations in developing aortic PVAT. **(A)** H&E staining of retroperitoneum *en bloc* from E18 and P3 CD1 mice (scale bar, 517.6 μm). **(B)** mRNA levels of indicated genes in thoracic from E18 and P3 CD1 mice (n=3 biological replicates of pooled aortas; mean+/−SEM) **(C)** UMAP projection of clustering of 17,957 cells from P3 thoracic aorta of P3 CD1 mice. **(D)** UMAP projections showing expression of smooth muscle marker genes (left) and immunostaining of PDGFRa (red), MYH11 (green), and DAPI (blue) in sections of E18 thoracic aorta (right) (scale bar, 271.8 μm). **(E)** UMAP projections showing expression of adipocyte marker genes (left) and immunostaining of PLIN1 (red) and DAPI (blue) in sections of P3 thoracic aorta. (scale bar, 543.5 μm). **(F)** UMAP projections showing expression of preadipocyte marker genes and immunostaining of PPARg (red), PDGFRa (green), and DAPI (blue) in sections of P3 thoracic aorta. Arrowheads in inset show preadipocytes (scale bar, 135.9 μm). **(G)** UMAP projection showing expression of intermediate cell marker genes and mRNA *in-situ* hybridization of *Bace2* (red), and DAPI (blue) in sections of P3 thoracic aorta. Arrowhead indicates intermediate cell. (scale bar, 145 μm). **(H)** UMAP projection showing expression of progenitor marker genes (left) and mRNA *in-situ* of hybridization of: (1) *Pi16* (green), and DAPI (blue) (middle); and (2) *Clec11a* (red), Pi16 (green), and DAPI (blue) (right) in P3 thoracic aorta. Arrowheads in [L] inset indicate progenitors. White arrowheads point to intermediate cells. Yellow arrowhead shows progenitor cell (right). (scale bar, 291 μm [L] and 145 μm [R]). **(I)** Model for thoracic aorta tissue organization at P3. PVAT; Perivascular Adipose Tissue; Lu; vessel lumen

To identify vascular adipocyte progenitor cells, we performed single cell RNA sequencing (scRNAseq) of all stromal-vascular cells isolated from the thoracic aorta at E18 and P3. We reasoned that these analyses would detect major shifts in the cellular composition of aortic tissue between E18 and P3, as the adipose tissue depot forms. Surprisingly, integration of the E18 and P3 scRNAseq datasets revealed nearly identical cell population structures (Fig. S1A-F) (Stuart et al., 2019). The only notable difference was the emergence of greater numbers of *Adipoq+* and *Adipoq+/Ucp1+* cells in the P3 dataset relative to E18 (Fig. S1F). We presume that the *Adipoq*-expressing cells present at E18 represent small nascent adipocytes that co-purified with stromal-vascular cells.

We focused our subsequent analysis on the P3 dataset given its greater representation of *Adipoq*/*Ucp1*+ adipocytes. Clustering analysis identified several expected cell populations, including endothelial cells, immune cells, mesothelium and neuronal cells (Fig. 2C, S1G). There was also a connected array of fibroblast cell clusters positioned between SMCs and adipocytes, including Intermediate Cells, Progenitors and Preadipocytes (Fig. 2C). Clustering analysis identified three groups of SMCs, all of which were marked by expression of the pan smooth muscle genes *Myh11* and *Acta2* (Fig 2D). MYH11+ SMCs were located around the aorta and did not express PDGFRa, a marker of fibroblast cells (Fig. 2D). Adipocytes were marked by specific expression of *Adipoq* and *Perilipin-1* (*Plin1*) (Fig. 2E). PLIN1-expressing adipocytes were exclusively located within adipose tissue lobes near the aorta (Fig 2E). Among the fibroblast groups, we identified a putative population of preadipocytes that clustered closest to adipocytes and expressed the master adipogenic regulator *Pparg* and the fibroblast marker *Pdgfra* (Fig. 2F). Co-staining of PPARg and PDGFRa in thoracic aorta revealed that preadipocytes were located at the periphery of the developing adipose tissue lobes (Fig. 2F). Another population of fibroblastic cells, which we termed intermediate cells, clustered closest to the SMCs and were marked by selective expression of *Bace2 and Clec11a* (Fig. 2G). Intermediate cells also expressed *Acta2*, albeit at lower levels than SMCs. *In situ* mRNA hybridization analysis revealed that *Bace2*-expressing cells were intimately associated with the outer wall of the aorta (Fig. 2G). The third fibroblast population, which we called progenitors, expressed enriched levels of *Pi16* and *Ly6a* (Fig S1D)*. Pi16* mRNA was localized in a majority of the adventitial cells surrounding the aorta. Co-staining of *Pi16* mRNA with the intermediate cell marker, *Clec11a,* revealed that progenitor cells were located more distally to the aorta than intermediate cells (Fig. 2H). Altogether, these analyses identified multiple fibroblast cell subtypes residing in discrete anatomic compartments within the developing aorta and its associated tissues (Fig. 2I).

### Isolation and characterization of adipogenic fibroblasts in perinatal thoracic aorta

We developed a fluorescence-activated cell sorting (FACS) strategy to isolate the three fibroblast populations (intermediate cells, progenitors, preadipocytes) and SMCs, based on their expression of cell surface markers. Dissociated cells from aortas were stained with CD31, CD45, and Ter119 to separate away endothelial, immune, and erythroid cells, respectively (Fig S2A). Progenitors were then identified and sorted based on high expression of LY6A (Fig. 3A, left panel, S2B). Preadipocytes were isolated from LY6A(−) cells based on the intermediate expression of CD142 and lack of CD200 expression (Fig. 3A, middle panel). SMCs were LY6A(−), CD200+ and CD142(−) (Fig. 3A, middle panel, S2C). Intermediate cells were purified from the CD200+;CD142+ cells by gating against CD317+ mesothelial cells (Fig. 3A, right). We performed bulk RNA sequencing on the four sorted cell populations to validate the sorting strategy and to obtain deep expression profiles for comprehensive pathway analyses. The cluster-defining genes identified from the single cell transcriptomes were mapped onto the sorted cell transcriptomes, revealing a high level of concordance between single cell and sorted cell gene expression profiles (Fig. 3B).

**Fig. 3.**
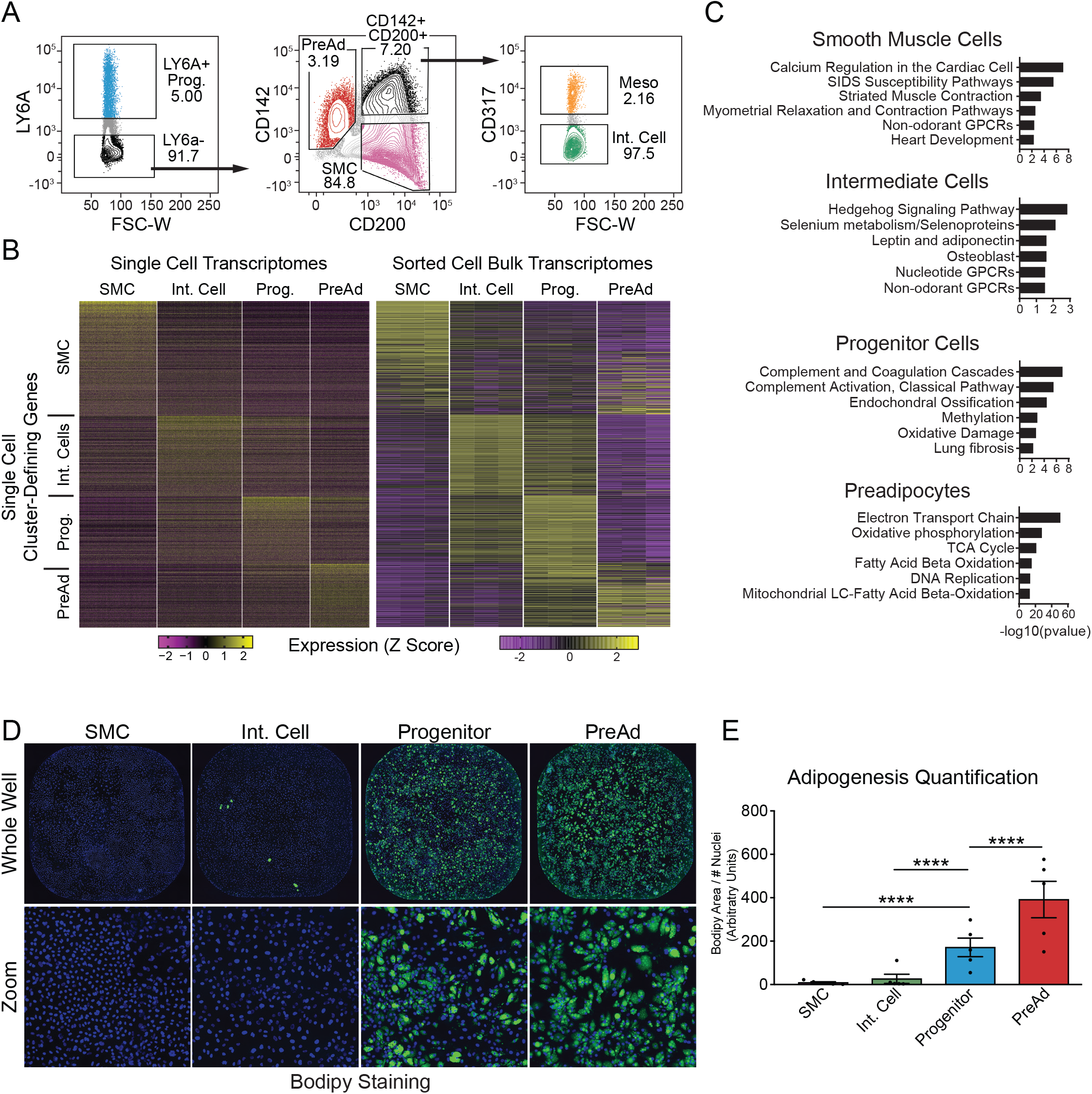
Isolation of adipogenic fibroblasts in perinatal thoracic aorta. **(A)** FACS isolation of fibroblast and smooth muscle cell populations. Live (FVS510−); Lin− (CD45−,CD31−,Ter119−) cells were gated as shown to isolate the following cell populations: Progenitors [LY6a high], Preadipocytes (PreAd) [LY6A−; CD142 intermediate; CD200−], Smooth Muscle Cells (SMC) [LY6A−; CD142−; CD200+], Intermediate Cells (Int) [LY6A−; CD142+; CD200+, CD317−]. Meso (Mesothelium) [LY6A−; CD142+; CD200+, CD317+]. (Representative plots from 10 separate experiments). **(B)** Expression heatmaps of Seurat-generated cluster-defining genes mapped on to sorted-cell RNA-Seq results (n=3 biological replicates per group). **(C)** Pathway analysis of cluster-defining genes for each cell population. Cluster-defining genes are defined as significantly differentially expressed genes with log2FC>1.5 in every pairwise comparison from sorted-cell bulk RNA-seq (n=3 biological replicates). **(D)** Bodipy (lipid; green) and Hoechst (DNA; blue) staining of indicated cell cultures following differentiation with adipogenic cocktail (Representative image of 5 separate experiments). **(E)** Quantification of adipocyte differentiation of cultures in [D] as Bodipy+ area above threshold/nuclear content (DAPI) (n=5 separate experiments; mean+/−SEM).

We compiled cell type-defining gene sets through differential gene expression analyses of the bulk RNAseq datasets; these stringent gene lists only comprised genes that were significantly and highly enriched in a cell type relative to all others (Fig. 3C). Pathway analyses of these gene lists uncovered a variety of cell type-enriched pathways. For example, “Calcium regulation” and “Striated Muscle contraction” were identified in SMCs. Intermediate cells were enriched for components of the “Hedgehog signaling pathway”. Progenitor cells demonstrated elevated expression of genes involved in “Complement Activation” as well as “Endochondral ossification”. Preadipocytes were notable for their enriched expression of genes related to “electron transport chain”, “oxidative phosphorylation”, and “TCA cycle”, corresponding to gene programs that are critical for the activity of thermogenic adipocytes.

To investigate the adipogenic potential of sorted cell populations, we performed *ex vivo* adipogenesis assays. Sorted cell populations were plated and treated with a standard adipogenic cocktail to stimulate adipocyte differentiation. SMCs and intermediate cells displayed almost no capacity to undergo adipocyte differentiation, as assayed by Bodipy staining for lipid accumulation (Fig. 3D,E). By contrast, progenitor cells and preadipocytes efficiently formed adipocytes, with the preadipocytes displaying the highest levels of adipogenesis (Fig. 3D,E). Overall, these studies provide a robust method for purification of specific aorta-associated cell populations and reveal a gradient of adipogenic competency from non-adipogenic SMCs to highly adipogenic preadipocytes.

### Adipogenic fibroblasts represent the major source of aortic PVAT in neonates

The contribution of smooth muscle and fibroblastic cells to perivascular adipocyte development was examined *in vivo* using genetic lineage tracing models (Fig. 4A). We used *Myh11-Cre* mice to investigate if SMCs develop into peri-aortic adipocytes. *Myh11* is a highly specific marker of SMCs and its expression was restricted to SMCs in our scRNAseq data set (Fig. S3A) (Miano et al., 1994). Analysis of *Myh11-Cre; tdTomato* reporter mice revealed uniform labeling of aortic smooth muscle. We did not detect any tdTomato+ adipocytes around the aorta, implying that SMCs do not give rise to adipocytes (Fig. 4B).

**Fig. 4.**
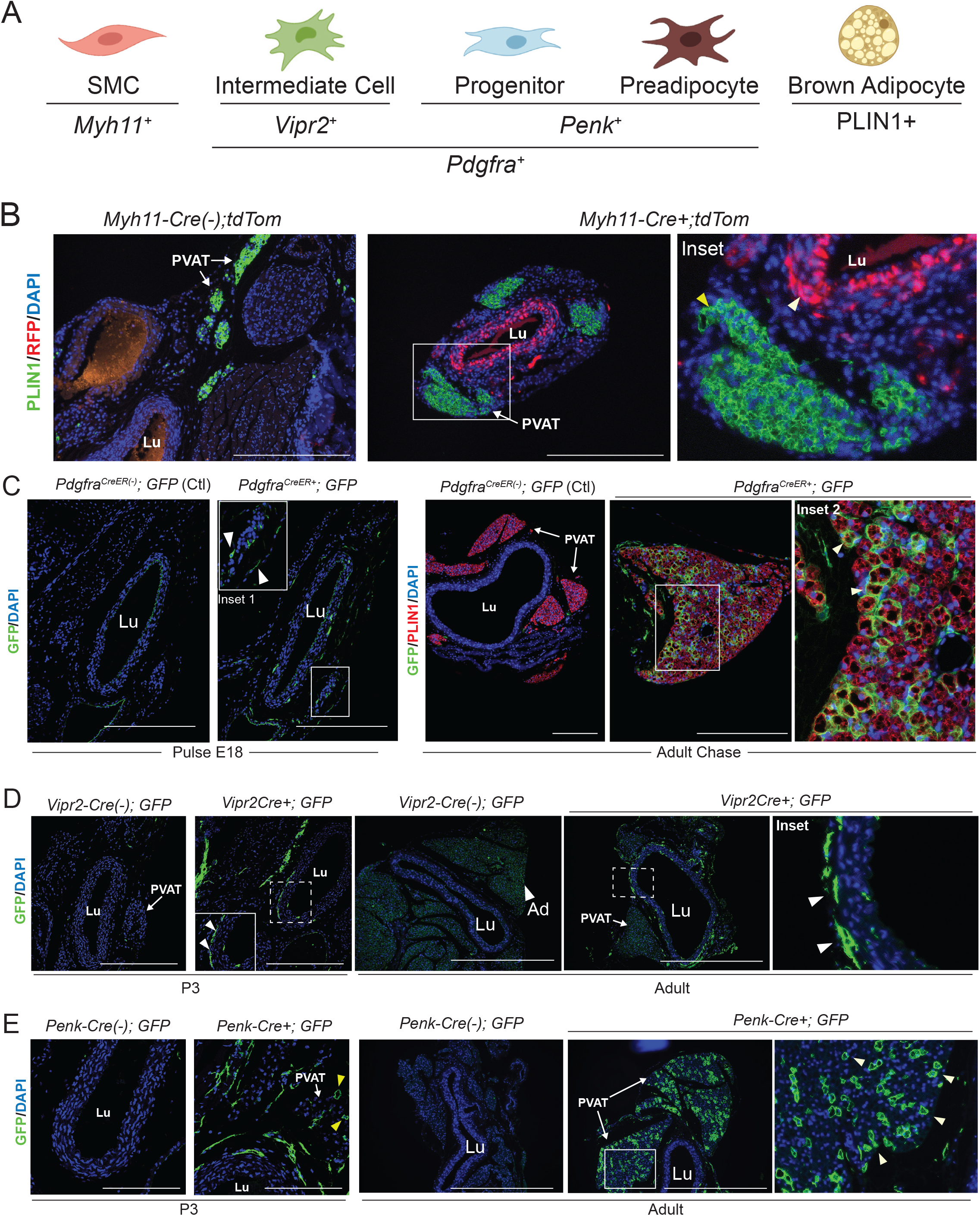
Adipogenic fibroblasts are the major source of aortic adipocytes in neonates. **(A)** Schematic of cell types in perinatal aorta and corresponding marker genes identified via RNA-Seq. **(B)** Immunostaining of PLIN1 (green), RFP (red) and DAPI (blue) in sections of P3 thoracic aorta from *Myh11-Cre(−); tdTom* (control) and *Myh11−Cre+; tdTom* reporter mice. White arrowhead in inset indicates labeled smooth muscle cells. Yellow arrowhead shows the absence of traced (RFP+) adipocytes. (n=2 *Cre−*; n=3 *Cre+*; scale bar, 271.8 μm). **(C)** Immunostaining of GFP (green), PLIN1 (red), and DAPI (blue) in sections of thoracic aorta from *Pdgfra-CreER(−); mTmG* (control) and *Pdgfra-CreER+; mTmG* mice harvested 24 hours (E18 Pulse) and 8 weeks (Adult Chase) following an E18 pulse with tamoxifen (100 mg/kg). Arrowhead in inset 1 indicates labeled fibroblasts. Arrowheads in inset 2 indicate GFP+ adipocytes. (Pulse: n=1 *Cre−*; n=3 *Cre+;* Chase: n=1 *Cre−*; n=2 *Cre+*; scale bar, 271.8 μm). **(D)** Immunostaining of GFP (green) and DAPI (blue) in sections of P3 and adult thoracic aorta from *Vipr2-Cre(−); mTmG* and *Vipr2-Cre+; mTmG* mice. Arrowheads indicate GFP expressing intermediate cells. (P3: n=3 *mTmG−*; n=4 *mTmG+;* Adult: n=1 *mTmG−*; n=4 *mTmG+*; scale bar, 290 μm (P3), 724.7 μm (adult)). **(E)** Immunostaining of GFP (green) and DAPI (blue) in sections of P3 and adult thoracic aorta from *Penk-Cre(−); mTmG* and *Penk-Cre+; mTmG+* mice. Yellow arrowheads indicate labeled fibroblasts. White arrowheads indicate labeled adipocytes. (P3: n=1 *Cre−*; n=3 *Cre+;* Adult: n=1 *Cre−*; n=3 *Cre+*; scale bar, 145um (P3), 724.7um (adult)). PVAT; Perivascular Adipose Tissue; Lu; vessel lumen

*Pdgfra* is expressed in all peri-aortic adventitial fibroblasts, including intermediate cells, progenitors and preadipocytes (Fig. S3A). To evaluate the contribution of these cell populations to adipocytes, we utilized tamoxifen-inducible *Pdgfra-CreER*; GFP reporter mice (*Pdgfra-CreER; mTmG)*. We administered one dose of tamoxifen to pregnant females on E18 to induce GFP expression in fibroblast cells. Analysis of embryos 24h later revealed extensive GFP expression in fibroblastic cells surrounding the aorta, including cells in the anatomic locations characteristic of intermediate cells, progenitors and preadipocytes (pulse) (Fig. 4C). In adult animals, we observed GFP expression in many mature adipocytes (marked by PLIN1-expression), demonstrating that perivascular adipocytes initially develop from an embryonically-specified *Pdgfra*+ fibroblastic lineage (Fig. 4C).

RNAseq analysis identified *Vipr2* as a selective marker gene of Intermediate cells relative to the other fibroblast subpopulations (Fig. S3B). We used *Vipr2*-*Cre;GFP* reporter mice to determine if intermediate cells develop into adipocytes. At P3, GFP expression was confined to fibroblastic cells associated with the outer SMC layer of the aorta, consistent with an intermediate cell identity. In adult mice, we observed a similar, robust pattern of GFP expression in fibroblast cells around the aorta, but an absence of GFP expression in adipocytes (Fig. 4D). Progenitors and preadipocytes were enriched for *Penk* expression, relative to other fibroblasts (Fig. S3A). In P3 *Penk-Cre:GFP* mice, GFP was expressed in peri-aortic fibroblastic cells, with GFP labeling of adipocytes becoming widespread in adulthood (Fig. 4E). Collectively, these experiments demonstrate that peri-aortic adipose tissue develops from fibroblastic progenitor cells, specifically progenitors and preadipocytes.

### Comparative analysis of thermogenic and white adipogenic progenitors

We next sought to determine whether perivascular progenitor cells were committed to the thermogenic lineage. We isolated the adipogenic populations from thoracic aorta (Progenitor and Preadipocyte) and preadipocytes from iWAT. These cell populations were then induced to undergo adipocyte differentiation in culture. The PVAT populations and iWAT cultures expressed comparable levels of general adipocyte marker genes (*Adipoq*, *Pparg2*), indicative of similar adipogenic competency (Fig. 5A, S4A). Notably, PVAT progenitors and preadipocytes but not iWAT preadipocytes adopted a thermogenic gene program (*Dio2*, *Ucp1*) following adipocyte differentiation (Fig. 5A, S4A).

**Fig. 5.**
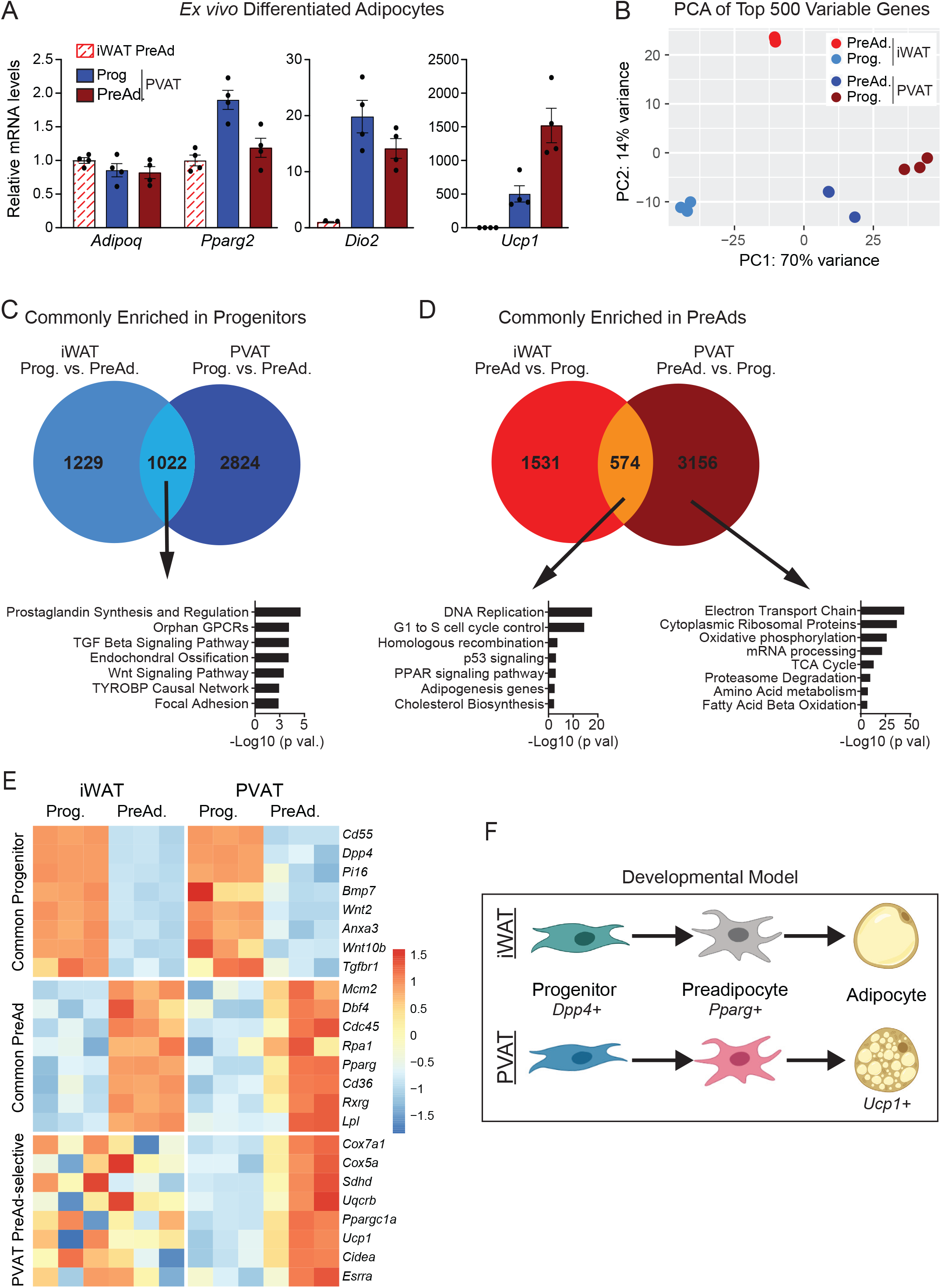
Comparative gene expression profiling of white and brown adipogenic cells. **(A)** mRNA levels of indicated genes in differentiated adipocytes derived from the following cell types of P3 CD1 mice: iWAT preadipocytes, thoracic perivascular progenitors and thoracic perivascular preadipocytes. (n=4 independent wells from pooled FACS populations per group; mean+/− SEM). **(B)** Principle component analysis (PCA) of the top 500 differentially expressed genes in sorted cell RNA Seq. (n=3 biological replicates per cell type). One PVAT PreAd datapoint includes two overlapping replicates. **(C)** Venn diagram of genes enriched in *progenitor populations* relative to *preadipocyte populations* from iWAT and PVAT. Pathway analysis of genes commonly enriched in both depots (significantly differentially expressed and LFC>0). **(D)** Venn diagram of genes enriched in *preadipocyte populations* relative to *progenitor populations*. Pathway analysis of genes commonly enriched in both depots (significantly differentially expressed and LFC>0) (left) and genes unique to PVAT (right). **(E)** Z-Score split heatmap of representative genes from GO analysis. Gene expression levels are calculated between cell types (i.e. PVAT Progenitors vs. PVAT PreAd) within a depot of origin (n=3 biological replicates per cell type). **(F)** Model for conserved ontogeny of adipose depot development.

Developing PVAT and iWAT contain analogous adipogenic populations: progenitor cells expressing genes such as *Pi16* and *Dpp4,* and preadipocytes expressing commitment markers *Pparg* and *Lpl.* To search for conserved gene programs controlling these two cellular states, we performed bulk RNA sequencing on freshly sorted progenitors and preadipocytes from aortic PVAT and iWAT. Principal component analysis showed that tissue origin accounted for much of the variance in the expression of the 500 most variable genes (Fig. 5B). We defined a consensus gene signature of progenitors, consisting of genes that were selectively expressed in progenitors vs. preadipocytes from both PVAT and iWAT (Fig. 5C,S4B). Pathway analysis of these genes identified enrichment of Prostaglandin synthesis, TGFβ and WNT signaling (Fig. 5C). The consensus preadipocyte genes were enriched for “PPAR signaling”, “adipogenesis” (*Pparg*, *Cd36*, and *Rxrg*), and an unexpected signature of cell cycle activity (*Mcm2*, *Cdc45*, and *Rpa1*) (Fig. 5D). Interestingly, PVAT preadipocytes specifically expressed genes related to electron transport chain, oxidative phosphorylation and the TCA cycle, including many genes important for the function (*Cox5a*, *Cox7a2*, and *Ucp1*) and transcriptional control (*Ppargc1a* and *Esrra*) of brown adipocytes (Fig. 5D,E). These data support a model in which uncommitted progenitor cells (enriched for expression of anti-adipogenic WNT and TGFβ pathways) develop into preadipocytes, with specific induction of a thermogenic gene program in PVAT preadipocytes (Fig. 5F).

### Identification of adipogenic smooth muscle cells in adult PVAT

To determine if the cell types observed in developing PVAT were also present in adults, we performed scRNAseq on the stromal-vascular fraction of thoracic aorta from 13-week-old mice. Analysis of the adult dataset alone revealed many of the adult cell groups were analogous to cell types observed in perinatal animals (Fig. 6A, S5A). Clustering analysis revealed three populations of adult fibroblastic cells (all *Pdgfra*^+^), two of which were present in the perinatal dataset: *Pi16*^+^/*Ly6a*^Hi^ progenitors and *Bace2*^+^/*Clec11a*^*Hi*^ intermediate cells (Fig. 6B). The third fibroblast population was defined by selective expression of *S100a4*, which we called “unknown fibroblasts”. *In situ* hybridization analysis of *Ly6a, Pi16* and *Bace2* showed that the spatial localization of intermediate and progenitor cells was conserved in adults, with intermediate cells located immediately adjacent to the aortic smooth muscle and progenitors located more distant radially (Fig. S5B,C). Of interest, we did not detect a fibroblastic preadipocyte (*Pparg*^+^/*Pdgfra*^+^) population. However, we observed a new cluster of SMCs (SMC 2) that expressed canonical smooth muscle markers *Myh11*, *Acta2* and *Tagln2*, along with *Pparg* and other adipocyte markers like *Lpl and Fabp4* (Fig. 6C).

**Fig. 6.**
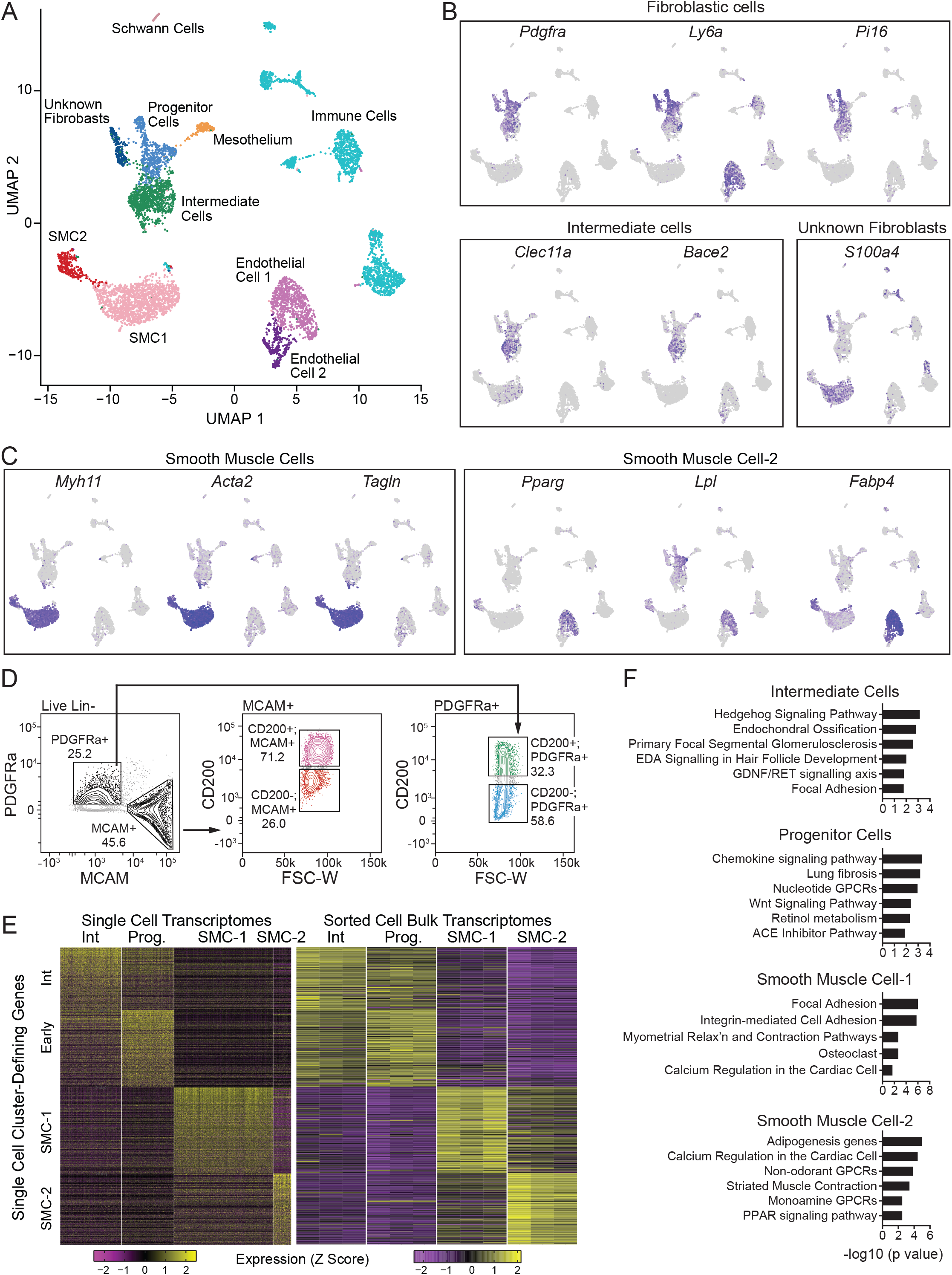
Identification of adipogenic smooth muscle cells in adult PVAT. **(A)** UMAP projection of clustering of 6,753 cells from adult thoracic aorta of pooled 13-week-old male CD1 mice. **(B)** UMAP projections showing expression of indicated fibroblast marker genes. **(C)** UMAP projections showing expression of indicated smooth muscle cell marker genes. **(D)** FACS isolation of fibroblast and smooth muscle cell populations. Live Lin(−) cells were gated to isolated the following cell populations: Intermediate Cells [PDGFRa+,MCAM−,CD200+], Progenitors [PDGFRa+,MCAM−,CD200−], Smooth Muscle Cells-1 (SMC-1) [PDGFRa−,MCAM+,CD200+], SMC-2 [PDGFRa−,MCAM+,CD200−]. (Representative image of 6 separate experiments). **(E)** Expression heatmap of Seurat-generated cluster defining genes mapped on to sorted-cell RNA-Seq results (n=3 biological replicates per group). **(F)** Pathway analysis of cluster-defining genes for each cell population. Cluster-defining genes were defined as significantly differentially expressed genes with log2FC>1.5 in every pairwise comparison (n=3 biological replicates).

We developed a FACS strategy to purify the following cell types from adult aortas for bulk RNAseq and functional characterization: (1) intermediate cells (PDGFRa+;CD200+); (2) progenitors and unknown fibroblasts (PDGFRa+;CD200−); SMC 1 (MCAM+; CD200+); and SMC-2 (MCAM+;CD200−) (Fig. 6D, S5D,E). The expression of cluster-defining genes from the single cell transcriptomes was mapped onto the sorted cell transcriptomes, revealing a high level of concordance (Fig. 6E, S5F,G). We determined cell type-defining gene sets by performing differential gene expression analysis and identifying genes that were enriched in each cell type compared to all others. This analysis again demonstrated that intermediate cells were enriched for components of the “Hedgehog signaling” pathway while progenitor cells were enriched for the “WNT signaling” pathway (Fig. 6F). SMC-1 cells were enriched for “focal adhesion” and “integrin-mediated cell adhesion pathways”. Finally, SMC-2 cells were enriched for “Adipogenesis genes”, “Calcium regulation in the cardiac cell”, and “PPAR signaling (Fig. 6F).

### Adipogenic activity of smooth muscle cells in adult PVAT

We tested the adipogenic differentiation competency of FACS-purified SMC and fibroblastic populations isolated from peri-aortic adipose tissue of adult mice. CD200+ SMCs (SMC-1) did not undergo adipogenesis, whereas CD200(−) SMCs (SMC-2), which expressed *Pparg* and other adipogenic genes, efficiently differentiated into lipid droplet-containing adipocytes (Fig. 7A). The adult fibroblast populations displayed similar adipogenic activity as their neonatal counterparts, with progenitor cells undergoing robust adipocyte differentiation, and intermediate cells displaying no adipogenic potential (Fig. 7A).

**Fig. 7.**
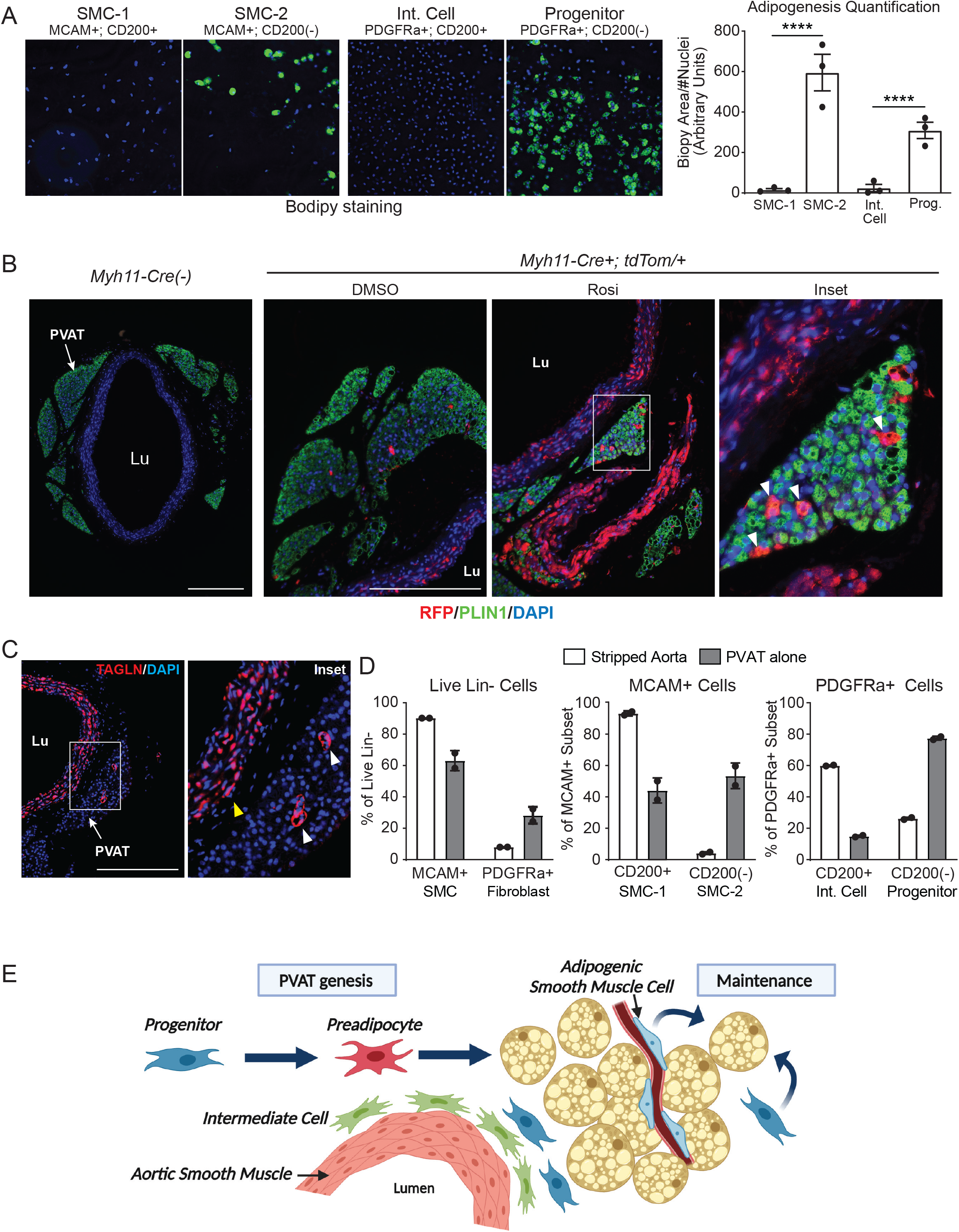
Adipogenic activity of smooth muscle cells in adult PVAT. **(A)** Cell populations were sort purified from adult aortas and induced to differentiate into adipocytes, followed by staining with Bodipy (lipid, green) and Hoechst (nuclei, blue) Quantification of adipocyte differentiation in cultures (right) (n=3 separate experiments; mean+/−SEM). **(B)** Immunostaining of RFP (red), PLIN1 (green), and DAPI (blue) in sections of thoracic aorta from adult *Myh11-Cre−; mTmG* and *Myh11-Cre+; mTmG* mice treated with vehicle control (DMSO) or rosiglitazone (Rosi). Arrowheads indicate mTomato-labeled adipocytes from *Myh11-Cre*+ cells. (n=1 *Cre−*; n=2 DMSO, n=2 Rosi; scale bar, 271.8 μm). **(C)** Immunostaining of TAGLN (red) and DAPI (blue) in sections of adult thoracic aorta. White arrowheads show adventitial smooth muscle cells. Yellow arrowhead indicates parenchymal smooth muscle cells of the aorta proper (scale bar, 271.8 μm). **(D)** Flow cytometry analysis of cell populations in stripped aorta or PVAT alone from adult CD-1 male mice (n=2 biological replicates of pooled aortas; mean+/−SEM). **(E)** Model of thoracic aorta adipose tissue organization PVAT; Perivascular Adipose Tissue; Lu; vessel lumen

To test if SMCs can contribute to perivascular adipocyte development in adult animals, we analyzed the aortas of adult *Myh11-Cre; tdTomato* reporter animals. Under baseline conditions, tdTomato+ adipocytes were detected, albeit at low levels, in adult peri-aortic fat. To further investigate the adipogenic activity of SMCs, we treated *Myh11-Cre; TdTomato* reporter mice with rosiglitazone to promote adipocyte turnover (Tang et al., 2011). Under these conditions, numerous, tdTomato+; PLIN1+ adipocytes were detected in thoracic PVAT, documenting the adipogenic capacity of SMCs in adult animals (Fig. 7B).

To test if fibroblastic cells are also competent to form adipocytes in adult PVAT, we tracked the fate of *Pdgfra*-expressing cells using *Pdgfra-CreER; GFP* reporter mice (Fig. S6A). These animals were treated with tamoxifen to induce GFP-expression in fibroblasts, followed by treatment with rosiglitazone. Immunofluorescence analysis revealed many GFP-marked adipocytes in aortic PVAT, indicating that fibroblastic cells can participate in *de novo* adipocyte formation (Fig. S6A). RNAseq analyses identified *Gli1* as a selective marker of intermediate cells vs. other fibroblast populations in adult aorta (Fig. S6B). To determine if intermediate cells can generate adipocytes, we performed lineage analysis with *Gli1-CreERT*;*GFP* mice (Fig. S6C). Upon tamoxifen administration, GFP expression was induced in intermediate fibroblasts lining the aortic smooth muscle and was not expressed in any adipocytes (Fig. S6C). Following rosiglitazone treatment of *Gli1-CreERT; GFP* mice, we did not detect GFP+ adipocytes, suggesting that intermediate cells are not a source of new adipocytes in adult animals (Fig. S6C).

Lastly, we investigated the anatomic localization of the two identified SMC subtypes in thoracic aortic tissues. Immunostaining for the pan SMC marker TAGLN identified SMC cells in the adipose tissue proper as well as in the aortic media around the lumen (Fig. 7C). Histologic analysis of the aorta in longitudinal section identified blood vessels coursing through the adipose tissue (Fig. S6D). To determine if the adipogenic *Pparg*^+^/CD200^−^ SMCs (SMC-2) were selectively enriched in the aortic adventitia, we separated adventitia/adipose tissue and aortic media, and isolated cells for flow cytometry analysis (Fig. 7D). The adventitia was highly enriched for CD200^−^ progenitors and the cleaned aorta was enriched for CD200^+^ intermediate cells, verifying the separation strategy. Notably, there was also striking enrichment of CD200- (adipogenic) SMCs (SMC-2) in the adventitia samples, whereas the cleaned aorta was enriched for CD200^+^ SMCs. These data suggest that SMCs within PVAT adopt an adipogenic profile and capacity in adult animals.

## Discussion

Aortic PVAT represents a unique type of thermogenic fat depot that is conserved in rodents and adult humans. This fat depot exhibits many of the defining properties of classical BAT, including high levels of UCP1 expression, responsiveness to cold exposure, dependence on the brown fat transcriptional factor EBF2, and resistance to obesity-induced inflammation. In this study, we integrated single cell transcriptomics, genetic fate mapping and *in vitro* differentiation assays to establish developmental hierarchies of thermogenic PVAT development and maintenance. We found that fibroblastic adipocyte progenitors seed the PVAT depot during the late fetal and early postnatal period. In contrast to conclusions from prior papers, our results suggest that *bona fide* SMCs do not significantly contribute to adipocyte formation during the genesis of PVAT. However, a new population of *Pparg*-expressing SMCs emerges in adult PVAT and this SMC subtype undergoes adipocyte differentiation *in vitro* and *in vivo* (Fig. 7E).

Aortic brown fat cells rapidly form and become organized into recognizable depots in the immediate postnatal period between E18 and P3. We hypothesized that this period would feature large shifts in progenitor cell phenotypes, correlating with the observed burst of adipogenesis. In actuality, we observed a striking similarity in the stromal cell composition of the depot immediately prior to birth (when no adipocytes are present) and after birth (when many adipocytes were present). This finding suggests that perivascular adipocyte progenitor cells are seeded earlier in development and are poised to undergo adipocyte differentiation in response to adipogenic cues. The window from E18 to P3 therefore provides a potentially powerful system to identify the signals that control adipogenesis *in vivo*.

Cell lineage tracking studies using the SMC-restricted *Myh11-Cre* driver indicates that differentiated SMCs do not contribute to aortic PVAT formation during the fetal and early postnatal period. This result contrasts with prior studies suggesting that perivascular adipocytes originate from SMCs, based on lineage tracing studies using *Tagln* (*Sm22*)-Cre (Chang et al., 2012, Ye et al., 2019). We have confirmed that *Tagln-Cre*+ cells give rise to nearly all perivascular structures, including aortic smooth muscle, adventitial fibroblasts, and adipocytes. We posit that *Tagln*-*Cre* is activated during early embryonic development in a progenitor cell population of undifferentiated cells (i.e. not SMCs) that do not express *Myh11* and are fated for a perivascular, but not necessarily SMC fate. Consistent with this notion, cells expressing *Tagln* at E8.5 develop into thermogenic adipocytes (Ye et al., 2019). Future studies will be needed to define the identity and developmental trajectories of this upstream early embryonic progenitor cell population.

We identified two distinct groups of adipogenic fibroblasts in fetal and early postnatal aortas, namely progenitors and preadipocytes. Progenitors were marked by expression of *Pi16* and *Dpp4*, whereas preadipocytes expressed certain adipocyte-related transcripts, including the master adipogenic transcription factor *Pparg*. Notably, analogous mesenchymal populations, expressing many of the same marker genes, were also recently identified in subcutaneous WAT (Merrick et al., 2019). In WAT, DPP4+ interstitial progenitor cells are fibroblasts which reside in the reticular interstitium and give rise to preadipocytes before finally differentiating into adipocytes (Merrick et al., 2019). Integrated transcriptomic analyses identified a common gene signature of progenitor cells in PVAT (i.e. progenitors) and WAT (interstitial progenitors). Pathway analysis identified enrichment of the anti-adipogenic WNT and TGFβ signaling pathways in both PVAT and WAT progenitors (Kennell and MacDougald, 2005; Longo et al., 2004; MacDougald and Mandrup, 2002). These pathways presumably play crucial roles in maintaining progenitor cell identity, at least in part, via suppressing adipogenic progression. We also identified a common gene signature of PVAT and WAT preadipocytes, which included many genes associated with adipogenesis, such as *Lpl* and *Cd36*. The common preadipocyte signature was also significantly enriched for genes linked to cell cycle progression. In this regard, adipocyte differentiation of preadipocyte cell lines requires a round of post-confluent mitosis (Tang and Lane, 2012; Tang et al., 2003). However, it remains unclear if the cell cycle gene profile identified in preadipocytes relates to cell cycle entry, exit or progression. Interestingly, many genes associated with fatty acid oxidation and mitochondrial biogenesis were selectively expressed in PVAT (brown) vs. WAT preadipocytes, indicating that metabolic specialization and divergence is already established in the preadipocyte state. Overall, these findings suggest that a conserved adipose cell lineage hierarchy operates in disparate fat depots, with less committed progenitor cells serving as a reservoir of preadipocytes.

We identified another interesting fibroblastic cell type in developing and adult aortas, which we termed intermediate cells due to their anatomic location in between the vascular smooth muscle cells and adventitial progenitors. Intermediate cells displayed selective enrichment of Hedgehog signaling genes. Previous studies identified a discrete population of sonic hedgehog-responsive cells located underneath the aorta, likely corresponding to intermediate cells (Passman et al., 2008). These cells did not undergo adipocyte differentiation either *in vitro* or *in vivo*. Understanding the physiologic function of intermediate cells remains a question of great interest. The intimate association of these cells with the outer layer of aortic smooth muscle, suggests a potential role in modulating vascular biology. In this regard, a recent report suggests that fibroblastic cells surrounding the vasculature have the capacity to differentiate into SMCs in the setting of dramatic vessel injury (Tang et al., 2020). Future work will determine if intermediate cells and/or other fibroblast subtypes adopt different phenotypes in pathologic states such as during atherogenesis or aneurysm development.

Our results suggest that adult PVAT does not retain a classical preadipocyte cell population, considered to be fibroblastic cells (i.e. *Pdgfra*+) that express adipogenic genes like *Pparg*. This result was surprising to us given that fibroblastic preadipocyte populations have been identified by us and others across several WAT depots. Rather, we found that adult PVAT contained a distinctive subset of SMCs that selectively expressed *Pparg* and other adipocyte-related genes. These *Pparg*+ SMCs clustered together with another SMC population, with both groups expressing similarly high levels of canonical SMC marker genes, including *Myh11*. Importantly, the *Pparg*^+^ SMC population displays robust potential to differentiate into adipocytes. Adult PVAT also contains progenitor cells, a fibroblast population with the potential to produce adipocytes. Our study suggests that both SMCs and progenitor cells contribute to the maintenance of adult PVAT. An elegant study by the Graff group revealed the existence of distinct lineages of embryonic and adult adipocyte progenitors in WAT (Jiang et al., 2014). Intriguingly, they found that adult adipocyte progenitors in WAT arise via an *Acta2*-expressing mural or SMC. It remains unclear if these *Acta2*+ cells in WAT or SMC in PVAT progress through a fibroblastic intermediate (i.e. preadipocyte) or differentiate directly into adipocytes. Another outstanding question is whether these different types of adipocyte progenitors in WAT and BAT/PVAT depots have the potential to differentiate into both white and brown or beige adipocytes.

In summary, this work defines the progenitor cells responsible for the development and maintenance of PVAT, a distinct thermogenic depot conserved in rodents and humans. A fibroblastic lineage, consisting of progenitor cells and preadipocytes, mediates the initial formation of aortic PVAT. After the postnatal period, this fat depot acquires an adipogenic subpopulation of SMCs. In adults, the aortic PVAT possesses two apparently independent sources of new adipocytes: adipogenic SMCs and fibroblastic progenitor cells. Additionally, a distinct fibroblast subtype is intimately associated with the aortic SMCs and does not contribute to adipocyte development or maintenance. The identification of the native progenitors for thermogenic adipocytes provides a critical framework for further investigations aimed at developing brown fat targeted therapeutics.

## Acknowledgements

We thank Drs. Xin Bi, Daniel Rader, Mitchell Lazar, Mark Kahn, Jorge-Henao-Mejia, and members of the Seale lab for thoughtful discussion. We thank Li Li and Min-Min Lu for additional tissue processing. We thank the University of Pennsylvania Diabetes Research Center for use of the Functional Genomics Core (P30-DK19525). This work was supported by NIH grants DK10300802 to P.S, DK120062 to A.R.A., and T32GM008216 to A.P.S.

## Author Contributions

A.R.A. A.P.S. and P.S were responsible for conceptualization, data analysis and writing/review. A.R.A. and A.P.S. contributed equally to the manuscript. A.R.A. and A.P.S. conducted the majority of the experiments. L.C. processed tissue sections and performed staining. C.O. assisted with experimental execution. R.S. performed cell capture and library preparation for perinatal single cell datasets. K.S. provided sequencing reagents and key experimental insight. K.B. assisted with data analysis.

## Lead Contact and Materials Availability

Further information and requests for resources and reagents should be directed to and will be fulfilled by the Lead Contact, Patrick Seale. Sequencing datasets will be available on GEO upon publication. All unique/stable reagents generated in this study are available from the Lead Contact without restriction.

## Methods

### Mice

Mice were housed under the care of University of Pennsylvania University Laboratory Animal Resources (ULAR), which provides both basic husbandry and veterinary care. Animals were raised at room temperature on standard chow with a 12-hour light/dark cycle. All mouse housing and husbandry occurred at RT (22°C) unless specified otherwise. For thermoneutral acclimation, 4- to 5-week old mice were housed at 30°C for one month. For chronic cold exposure, 8- to 10-week-old mice were pair-housed in cages at 4°C for one week. Mice were fed a western diet composed of 40% fat and 0.15% cholesterol from Research Diets (D12079B). Tamoxifen (Sigma [T5648], stock 20 mg/mL in corn oil) was intraperitoneally injected at a dose of 100 mg/kg for one day or five consecutive days. Mice were treated with rosiglitazone (250 nMoles/d) (Cayman catalog #11884) for the experimental time course indicated in text. Experiments performed on embryonic and perinatal mice were conducted on male and female mice. Experiments at adult time points were performed in male and female mice between the ages of 8 to 12 weeks at the onset of the experiment.

*C57/Bl6N* (RRID:IMSR_TAC:B6) mice were obtained from Taconic. CD1 (RRID:IMSR_CRL:024) mice were obtained from Charles River. The following strains were obtained from the Jackson Laboratory: *mTmG* (strain name: B6.129(Cg)- *Gt(ROSA)26Sor*^*tm4(ACTB-tdTomato,-EGFP)Luo*^/J, RRID:IMSR_JAX:007676), *tdTom* (strain name: B6.Cg-*Gt(ROSA)26Sor*^*tm14(CAG-tdTomato)Hze*^/J, RRID:IMSR_JAX:007914), *Myh11-Cre* (strain name: B6.Cg-Tg(Myh11-cre,-EGFP)2Mik/J, RRID:IMSR_JAX:007742), *Pdgfra-CreER* (strain name: B6.129S-*Pdgfra*^*tm1.1(cre/ERT2)Blh*^/J, RRID:IMSR_JAX:032770), *Gli1-CreER* (strain name: *Gli1*^*tm3(cre/ERT2)Alj*^/J, RRID:IMSR_JAX:007913), *Penk-Cre* (strain name: B6;129S-*Penk*^*tm2(cre)Hze*^/J, RRID:IMSR_JAX:025112), and *Vipr2-Cre* (strain name: B6.Cg-*Vipr2*^*em1.1(cre)Hze*^/J, RRID:IMSR_JAX:031332). The following stock was generated by Patrick Seale: *AdipoqCre*; *Ebf2 loxP/loxP* (Angueira et al., 2020).

### Histology and Immunofluorescence

Tissues were fixed in 4% PFA overnight, washed in PBS, dehydrated in ethanol, paraffin-embedded and sectioned. For *en bloc* retroperitoneal sections, perinatal mice were euthanized by decapitation, the thoracic viscera were removed, and the tissue was placed in 4% PFA overnight. Adult mice were perfused with 5 mL of 4% PFA and tissues were harvested as above. Slides were deparaffinized and heat-antigen retrieved in Bulls Eye Decloaking buffer (Biocare). Slides were incubated in indicated primary antibodies overnight, stained with secondary antibody, and developed with Tyramide Signal Amplification (TSA, Akoya Biosciences). Primary antibodies used for staining were: PERILIPIN: (1:200, CST: 3470, RRID:AB_2167268), PDGFRA: (1:50, Novus Biologicals: AF1062, RRID:AB_2236897), MYH11: (1:100, Abcam: ab53219, RRID:AB_2147146), PPARG: (1:500, Invitrogen: MA5-14889, RRID:AB_10985650), RFP: (1:250, VWR Scientific, 600-401-379, RRID:AB_2209751), GFP (1:500, Abcam: AB6673, RRID:AB_305643), CD200: (1:25, Novus Biologicals: AF2724, RRID:AB_416669), ACTA2: Sigma: (1:200, A2547, RRID:AB_476701), TAGLN: (1:100, Abcam: ab10135, RRID:AB_2255631), and DAPI (1:1000 Roche). RNA *in situ* hybridizations were performed using the RNAscope system (Advanced Cell Diagnostics; 323100, 323120) with the following probes: *Bace2* (407151), *Clec11a* (583301), *Pi16* (451311-C2), and *Ly6a* (427571-C2). Images were captured on a Leica TCS SP8 confocal microscope or Keyence inverted microscope.

### RNA Extraction, qRT-PCR and RNA Sequencing analysis

Total RNA was extracted using TRIzol (Invitrogen) and Purelink RNA columns (Fisher). For processing of RNA from cultured primary adipocytes, RNA was extracted using Trizol (Invitrogen) combined with PicoPure RNA Isolation Kit (Fisher). mRNA was reverse transcribed into cDNA using the ABI High-Capacity cDNA Synthesis kit (ABI). Real-time PCR was performed on an ABI7900HT PCR machine using SYBR green fluorescent dye (Applied Biosystems). Fold changes were calculated using the ΔΔCT method, with *Tata Binding Protein* (*Tbp*) mRNA serving as a normalization control. RNA sequencing from sorted progenitors was performed at Genewiz with the following procedure. RNA was extracted using Trizol LS (Invitrogen) and quantified using Qubit and TapeStation. RNA sequencing libraries were made using the SMART-Seq v4 Ultra Low Input Kit. Samples were sequenced on an Illumina HiSeq with a 2×150 Paired End (PE) configuration.

FASTQ files were aligned to mm10 using STAR (2.5.2a) with quantMode = GeneCounts. The unstranded genecounts for each sample were analyzed for differential gene expression using the R (3.6.3) package DESeq2 (1.26.0). In general, for each experiment involving an RNA seq dataset, the DESeq2 model was built with the design (~CellType) where “CellType” refers to the different sorted populations, with all CellTypes included in the model. Pairwise differential expression was calculated using the DESeq2 function “results” with thresholds of alpha=0.1. Log2 Fold change thresholds were used as indicated. ENSEMBLID’s were converted to gene symbols using the R package org.Mm.eg.db (3.10.0). Pathway analysis was performed via WikiPathways with annotations for Mus Musculus using the package clusterProfiler (3.13.3) and rWikiPathways (1.6.1) (Figures 4 and 6). Pathway analysis was performed using Enrichr WikiPathways 2019 Mouse for Figure 3.

Heatmaps represent either Log2 fold change versus row mean or row score as indicated as indicated. Underlying data for heatmaps came from the DESeq2 regularized log transform of the normalized count values using the DESeq2 function “rlog”. Figures showing side by side scRNA-seq and bulk RNA seq heatmaps contain the exact same genes in the same order on both heatmaps.

### scRNA Seq

Single cells were isolated from dissected aortas and flow sorted to isolate live (DAPI−) cells and remove debris. Cells were collected as either CD45+ (immune) and CD45− (non-immune) cells. For adult single cell RNA Sequencing, CD45+ cells were mixed with CD45− cells to achieve a proportion of ~20%. Single cell transcriptomes were collected as described previously using the 10X Genomics platform with v2 (perinatal) and v3 (adult) chemistry (10× Genomics, Pleasanton, CA) (Merrick et al., 2019). The libraries were sequenced using an Illumina HiSeq 4000. The reads were processed using the CellRanger pipeline and R Studio using Seuratv3 (Merrick et al., 2019, Stuart et al., 2019). Cells were filtered based on their expression of a minimum number of genes (E18/P3: 500 < nFeatureRNA < 3000; Adult: 250 < nFeature RNA < 4000) and mitochondrial genome (<10%) abundance for each dataset individually. Regression was performed using ScaleData function for the following variables: percentage of mitochondrial reads and cell cycle phase (G2M.Score/S.Score). Dimensionality reduction was performed in Seurat using UMAP and differential gene expression between clusters was performed using FindMarkers function. For all datasets, immune cells were collapsed into one cluster for simplicity. For the perinatal dataset, two clusters of mesothelial cells were collapsed to generate one mesothelial cluster. For the adult dataset, initial clustering revealed that SMC population 1 separated into two clusters, but the effect size of gene expression between the two groups was small, so they were collapsed to form SMC population 1. Feature-plots, dot-plots and heat-maps were generated using Seurat.

### E18/P3 combined analysis

Dataset integration was performed using the Seurat functions FindIntegrationAnchors followed by IntegrateData. Scaling and regression were performed using the Seurat ScaleData function for the following variables: percentage of mitochondrial reads and cell cycle phase (G2M.Score/S.Score). Dimensionality reduction was performed in Seurat using UMAP and differential gene expression between clusters was performed using FindMarkers function. For all datasets, immune cells were collapsed into one cluster for simplicity. Single gene UMAP feature plots showing expression of select genes at E18 and P3 were generated for the RNA assay using a modified version of the Seurat FeaturePlot function. (This modified version plots both the E18 and P3 UMAPs on the same color/expression scale, while the default Seurat function does not).

### Generation of single cell suspensions from aorta

Aorta were isolated from mice, minced and placed into digestion medium (DMEM, Collagenase D: 6.1mg/ml (Roche), Dispase II: 2.4 mg/ml (Roche) and placed at 37°C with constant agitation at 200 rpm. To enrich for cell populations with differential sensitivity to digestion, we utilized the following procedures. For embryonic/newborn mice, 20% of the digestion was quenched at 15 minutes, 20% at 20 minutes and the remaining 60% at 25 minutes (embryos/perinatal). For adult aortas 50% of the digestion was quenched at 30 minutes, 50% at 60 minutes. For stripped aorta flow cytometry analysis, perivascular adipose tissue was removed from the aorta and digested separately from the cleaned aorta. Tissue digestions were quenched with an equal volume of complete medium (DMEM/10% FBS). Dissociated cells were suspended using a P1000 pipette and filtered through a 100 μm filter. Cells were then pelleted at 400 g for 4 min and RBCs were lysed in 155mM NH_4_Cl, 12mM NaHCO_3_, and 0.1mM EDTA for 4 min. An equal volume of complete medium was added and the cells were filtered through a 40um filter for downstream analyses.

### Flow Cytometry and Sorting

Isolated cells were suspended in FACS buffer (HBSS, 3% 0.45uM filtered FBS; Fischer). Cells were stained in FACS buffer with antibodies (see below) for 1 hr at 4°C in the dark. Cells were washed 2x with cold FACS buffer and sorted on a BD FACS Aria cell sorter (BD Bioscences) with a 100 μm nozzle as previously described (Merrick et al., 2019). All compensation was performed at the time of acquisition in BD FACS DIVA software using single stained cells or compensation beads (BioLegend catalog no. A10497). The following antibodies were used for FACS protocols. **P3 Aorta:** LY6A-BV711 (1:100 Biolegend-108131 RRID:AB_2562241),CD317-PerCp-Cy5.5 (1:100 Biolegend-127022 RRID:AB_2566647), CD200-AF488 (1:100 Biorad-MCA1958A488), CD45-APC/Cy7 (1:1000, Biolegend-103115 RRID:AB_312980), CD31-APC/Fire750 (1:1000 Biolegend-102528 RRID:AB_2721491), Ter119-APC/Cy7 (1:1000 Biolegend 116223 RRID_AB:2137788), CD142-CF647 (1:1000 Sino Biological), FVS510 (1:1000 BD Biosciences 654406). **P3 iWAT:** CD45-APC/Cy7 (1:1000, Biolegend-103115 RRID:AB_312980), CD31-APC/Fire750 (1:1000 Biolegend-102528 RRID:AB_2721491), CD142-CF647 (1:1000 Sino Biological), CD26(DPP4)-FITC (Biolegend, San Diego, CA, cat# 302704 1:100); ICAM1-PE/Cy7 (Biolegend cat# 353116 1:100). **Adult Aorta:** CD200-AF488 (1:100 Biorad-MCA1958A488), CD45-APC/Cy7 (1:1000, Biolegend-103115 RRID:AB_312980), CD31-APC/Fire750 (1:1000 Biolegend-102528 RRID:AB_2721491), Ter119-APC/Cy7 (1:1000 Biolegend 116223 RRID_AB:2137788), MCAM-PerCP/Cy5.5 (1:100 Biolegend 134709 RRID:AB_11204083), PDGFRa-PE/Cy7 (1:200 Biolegend 135912 RRID:AB_2715974), and DAPI (1:10000 Roche 10236276001).

### Cell Culture

Sort-purified cells were plated on 384-well Cellbind plates (Sigma-Aldrich catalog no. CLS3770) and cultured in DMEM/F12 supplemented with 10% FBS and Primocin (50 μg/mL). Cells were plated at a density of 10,000-30,000 cells per well and induced to differentiate into adipocytes within 3 days of initial plating. Cell density and time until differentiation was kept consistent within each experiment. Adipogenic differentiation cocktail consisted of 1 μM dexamethasone (Sigma catalog no. D4902), 0.5 μM isobutylmethylxanthine (Sigma catalog no. I7018), 125 nM indomethacin, 20 nM insulin, and 1 nM T3 for 48 hours. Following two days of induction, media was changed to 20 nM insulin and 1 nM T3. Medium was changed every two days for six more days.

### Adipogenesis Assays and Quantification

Adipogenesis was assessed by staining with Bodipy 493/503 (Invitrogen catalog no. D3922) for lipid accumulation and Hoechst 33342 (Thermo Fisher catalog no. 62249) for nuclei, as described previously (Merrick et al., 2019). Briefly, cells were differentiated in 384 well tissue culture plates (Sigma-Aldrich catalog no. CLS3770), fixed with 4% paraformaldehyde, stained, and imaged on a Keyence inverted microscope (BZX-710) with the following filters: DAPI (ex, 360/40 nm; em, 460/50 nm; Keyence, OP-87762) and GFP (ex, 470/40 nm; em, 525/50 nm; Keyence, OP-87763) filters. Images were acquired at 20x in a 7×7 tiled grid and stitched to capture the entirety of each well. Tiling and stitching were performed with Keyence BZ-X Viewer software. Image quantification was performed automatically in ImageJ using a macro which: 1) Split images into component channels, 2a) for the nuclei channel applied a 3-Sigma Gaussian blur, performed thresholding to identify signal above background, performed watershed to segmentation, and counted the number of nuclei, 2b) for the lipid channel applied a 2-Sigma Gaussian blurm performed thresholding to identify signal above background, and counted the area (#of pixels) with signal above threshold. The amount of adipogenesis was calculated as Lipid Area/#nuclei.

### Statistical Analysis

No power calculations were performed prior to initiation of the study. No mice were omitted from the study. All individual data points were plotted to assay normality. Experiments from a common pool of digested tissue or independent pools processed separately are indicated in the text. Two sample t tests were performed where comparisons between two groups were being assayed. One way ANOVAs with pairwise comparisons corrected with the Holm Sidak method were used for comparisons between more than two groups. Two way ANOVAs with multiple comparisons corrected with the Holm Sidak method were performed for adipogenesis experiments (within each experimental day compare between cell types). # indicates p value < 0.1; * indicates p value < 0.05, ** indicates p value < 0.01, *** indicates p value < 0.001, **** indicates p value < 0.0001.

---

**Fig. S1:**
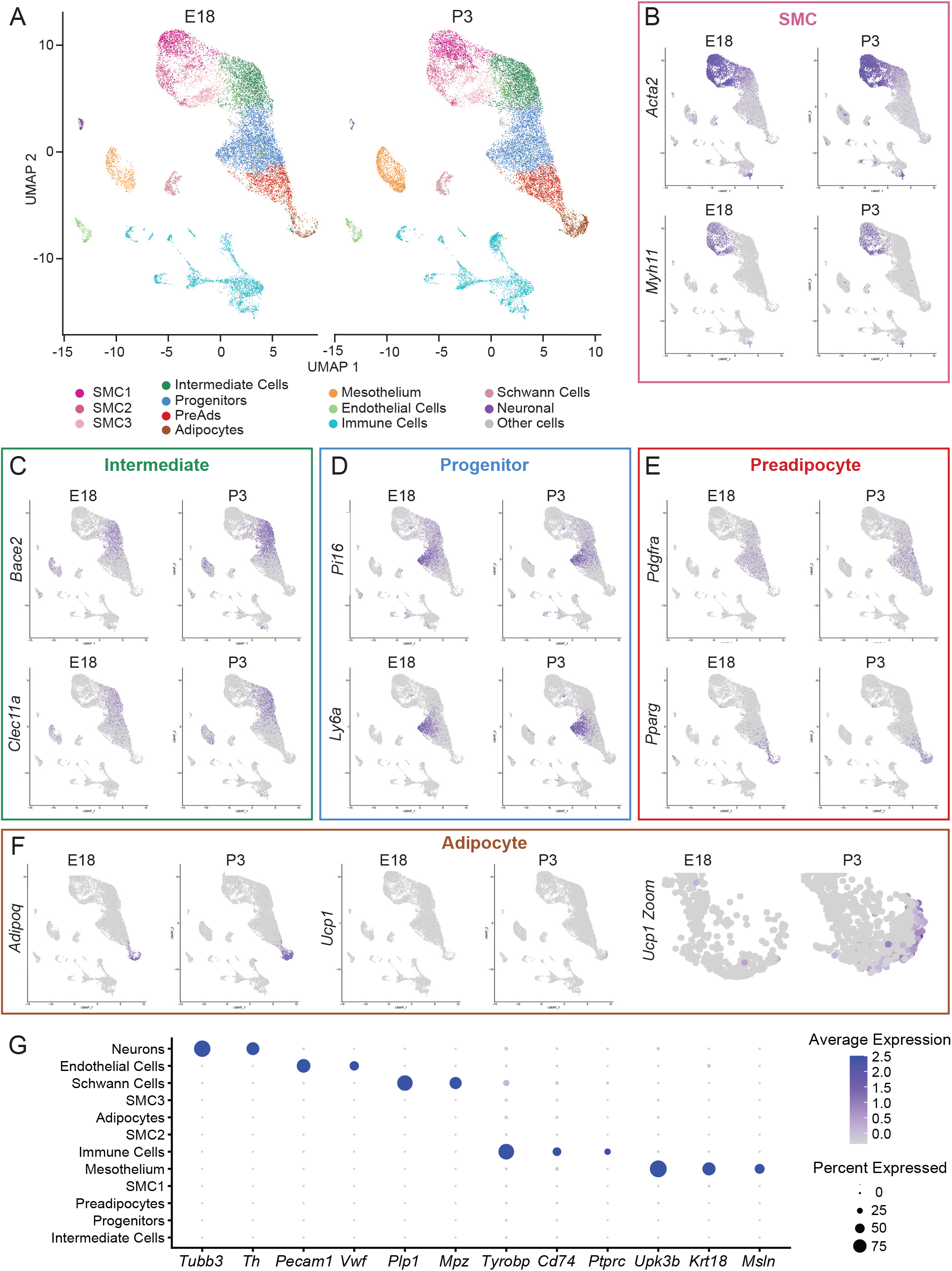
Single cell profiling analyses of aortas from E18 and P3 mice (Related to Fig. 2) **(A)** UMAP projection of clustering of cells from E18 and P3 thoracic aorta of pooled CD1 mice. **(B-F)** UMAP projections showing expression of indicated genes for: smooth muscle cells (SMC) (B); Intermediate cells (C); Progenitors (D); Preadipocytes (PreAds) (E); and Adipocytes (F). **(G)** Expression dotplot of indicated genes for cell clusters for P3 aorta.

**Fig. S2:**
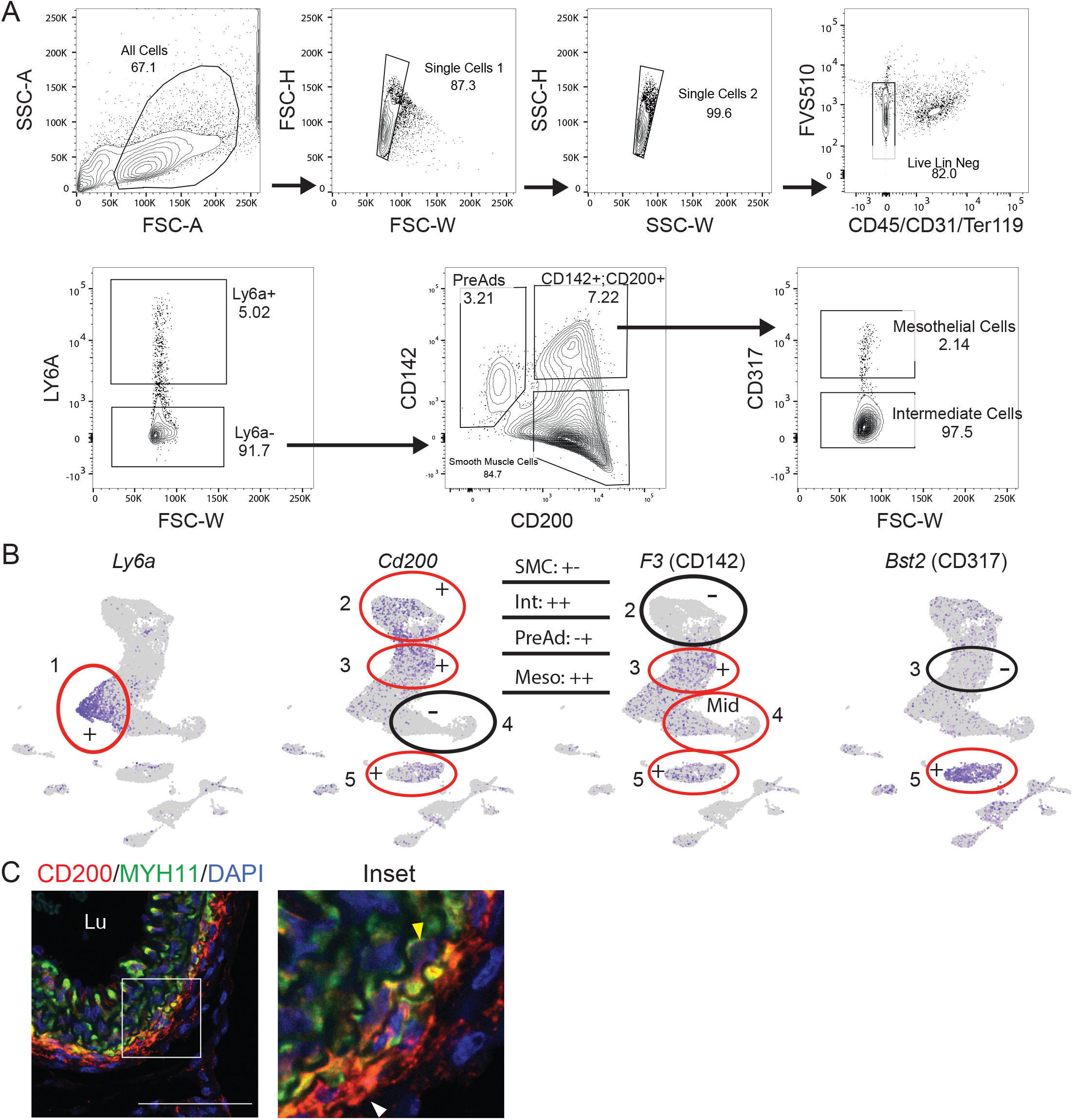
Purification of smooth muscle and fibroblastic cells from thoracic aortas (Related to Fig. 3) **(A)** FACS isolation of fibroblastic and smooth muscle populations. Dissociated cells were: (1) gated on SSC-A and FSC-A to exclude debris; (2) FSC-H vs. FSC-W then SSC-H vs. SSC-W to isolate single cells; and (3) gated on Live (FVS510−) Lin− (CD45−,CD31−,Ter119−) cells. Depicted sort gates were used to isolate the following cell populations: Progenitors [LY6A high], Preadipocytes (PreAd) [LY6A−, CD142 mid; CD200-]; Smooth Muscle Cells (SMC) [LY6A−; CD142−; CD200+], Intermediate Cells (Int) [LY6A−; CD142+; CD200+, CD317−]. Meso (Mesothelium) [LY6A−; CD142+; CD200+, CD317+]. (Representative image of 10 expts). **(B)** UMAP projection of P3 thoracic aorta single-cell data showing expression of indicated genes used in the above sorting strategy. Red circles represent presence of expression. Black circles indicate lack of expression. Circles correspond to the following clusters from Fig 2: 1- Progenitor Cells, 2-SMCs, 3-Intermediate Cells, 4 Preadipocytes and Adipocytes, 5-Mesothelium. **(C)** Immunostaining of CD200 (red), MYH11 (green), and DAPI (blue) in sections of P3 thoracic aorta. White arrowhead shows an Intermediate cell. Yellow arrowhead shows an SMC (scale bar, 65um; Lu: lumen).

**Fig. S3:**
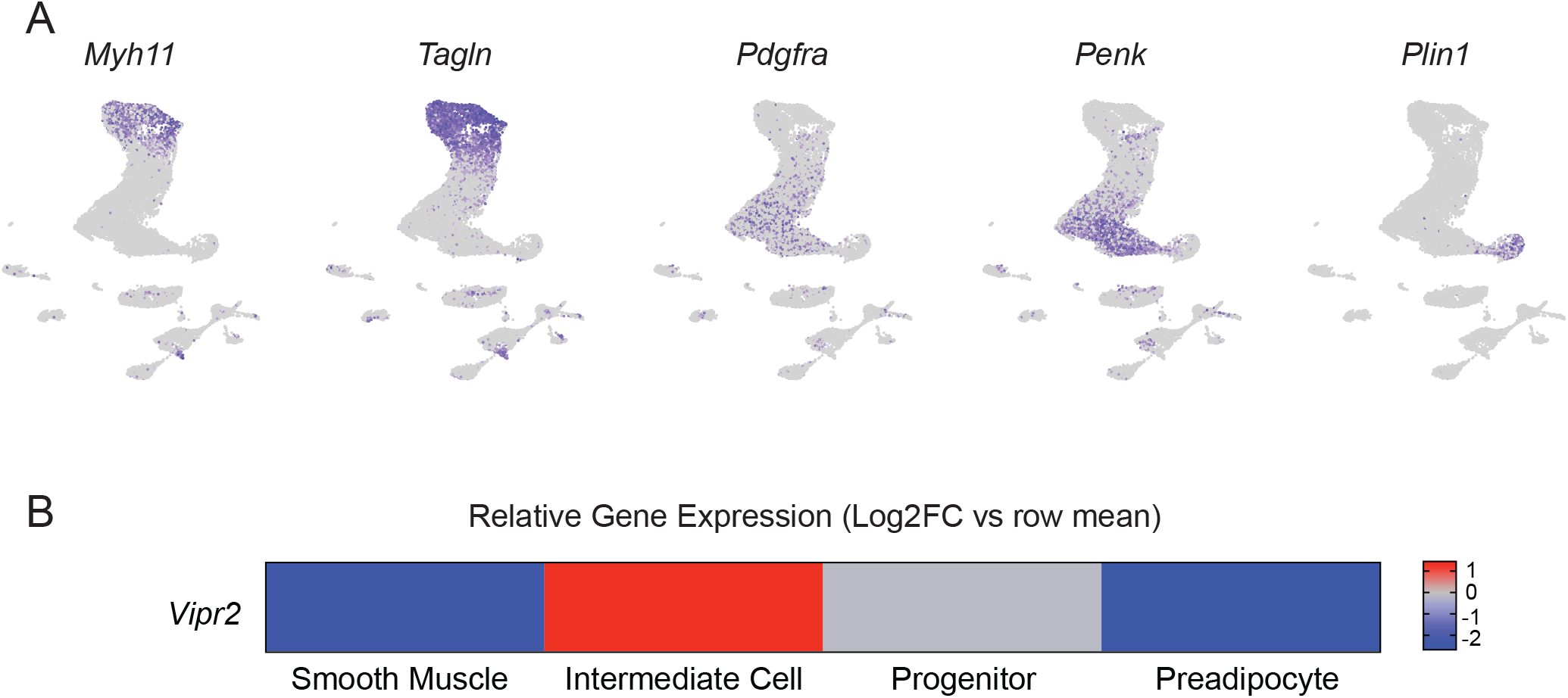
Expression profiles of Cre-driver mouse lines (Related to Fig. 4) (A) UMAP projection showing expression of indicated genes used for Cre and CreER driver mouse strains. (B) Log2FC expression analyses of Vipr in indicated cell types from sorted cell bulk RNAseq datasets.

**Fig. S4:**
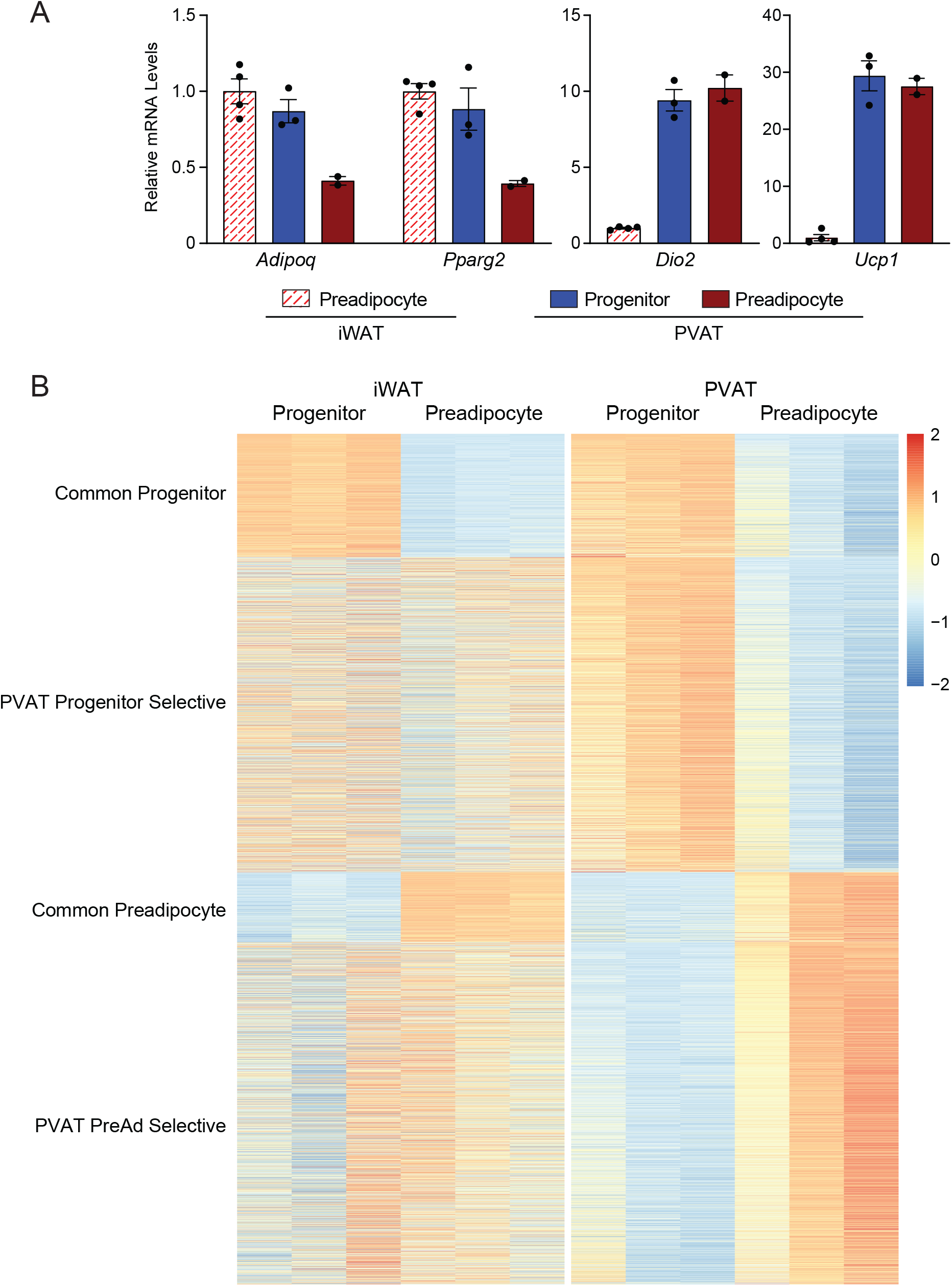
Gene profiling analyses of iWAT vs. PVAT progenitors and preadipocytes (Related to Fig. 5) **(A)** Relative mRNA levels of indicated genes in differentiated primary adipocytes derived from: iWAT preadipocytes, thoracic aortic PVAT progenitors. thoracic aortic PVAT preadipocyte cells from P3 CD1 mice. Second experimental replicate performed on different day (n=2-4 individual wells from pooled sort per group; mean+/− SEM). **(B)** Z-Score split heatmap of of genes from overlaps depicted in Fig. 4C,D. Gene expression levels are calculated between cell types (i.e. PVAT Progenitors vs PVAT PreAd) within a tissue of origin (n=3 biological replicates per cell type).

**Fig. S5:**
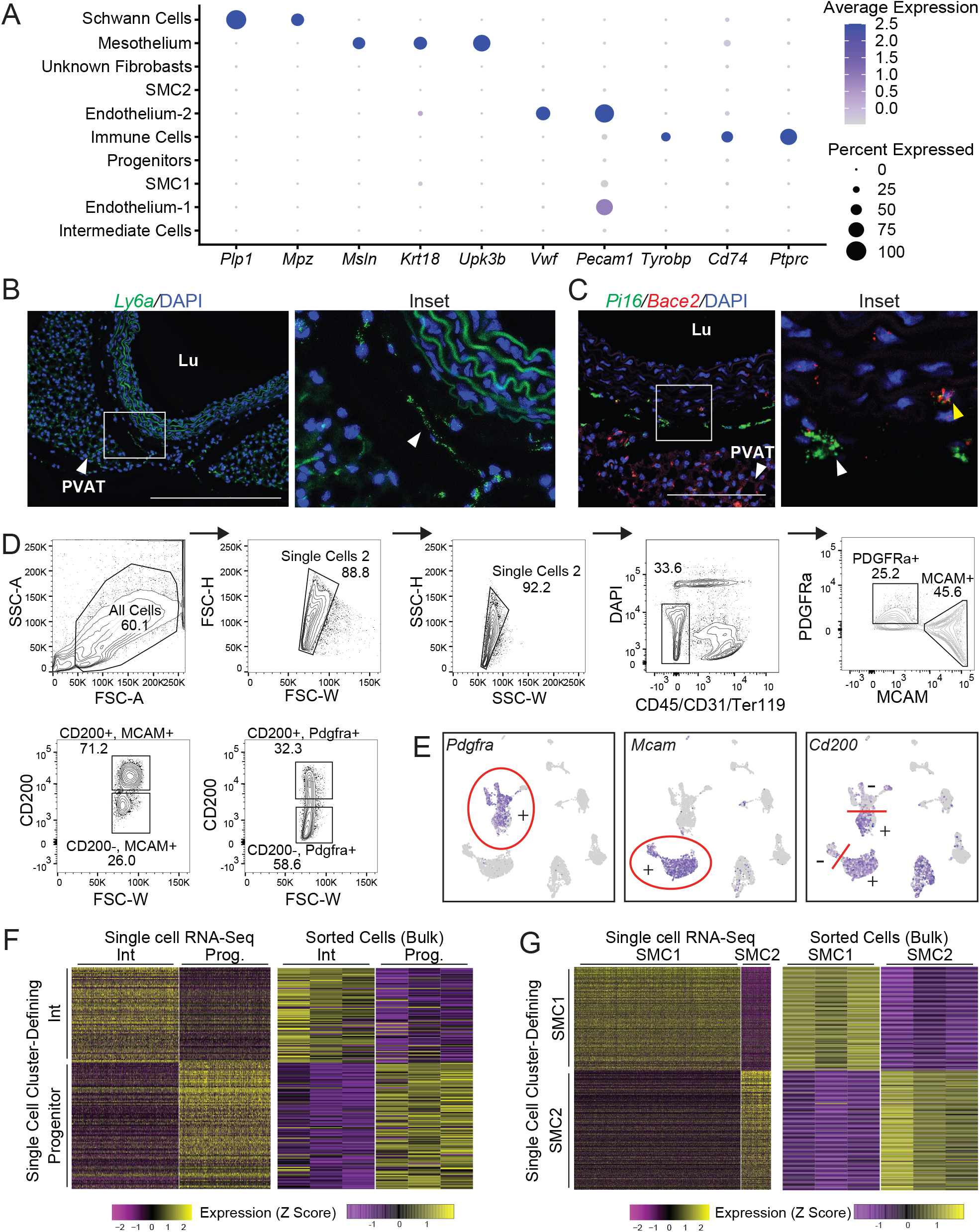
Purification and analysis of aorta-associated cells from adult animals (Related to Fig. 6) **(A)** Expression dot plot of indicated genes for cell clusters from adult thoracic aorta. **(B)** mRNA *in situ* hybridization of *Ly6a* (green) in adult thoracic aorta. Arrowhead in inset shows progenitor cell. DAPI (blue) stains nuclei. (scale bar, 362.3 μm). Lu: aorta lumen. **(C)** mRNA *in situ* hybridization of *Pi16* (green) and *Bace2* (red). White arrowhead shows progenitor cell. Yellow arrowhead shows intermediate cell. (scale bar, 145um). **(D)** FACS isolation of fibroblasts and smooth muscle cells (SMCs). To exclude debris and isolate single live cells, we gated on: (1) SSC-A and FSC-A; (2) FSC-H vs FSC-W; and (3) SSC-H vs SSC-W. Live (DAPI−); Lin− (CD45−,CD31−,Ter119−) cells were then selected. Further selection was then performed to isolate: Intermediate Cells [PDGFRa+,MCAM−,CD200+], Progenitors [PDGFRa+,MCAM−,CD200−], SMC1 [PDGFRa−,MCAM+,CD200+], SMC2 [PDGFRA−,MCAM+,CD200−]. (Representative image of 6 separate experiments). **(E)** UMAP projections showing expression of indicated genes used for sorting strategy. **(F,G)** Expression heatmap of Seurat-generated cluster-defining genes mapped on to sorted-cell RNA-Seq results for fibroblast (F) and SMC populations (n=3).

**Fig. S6:**
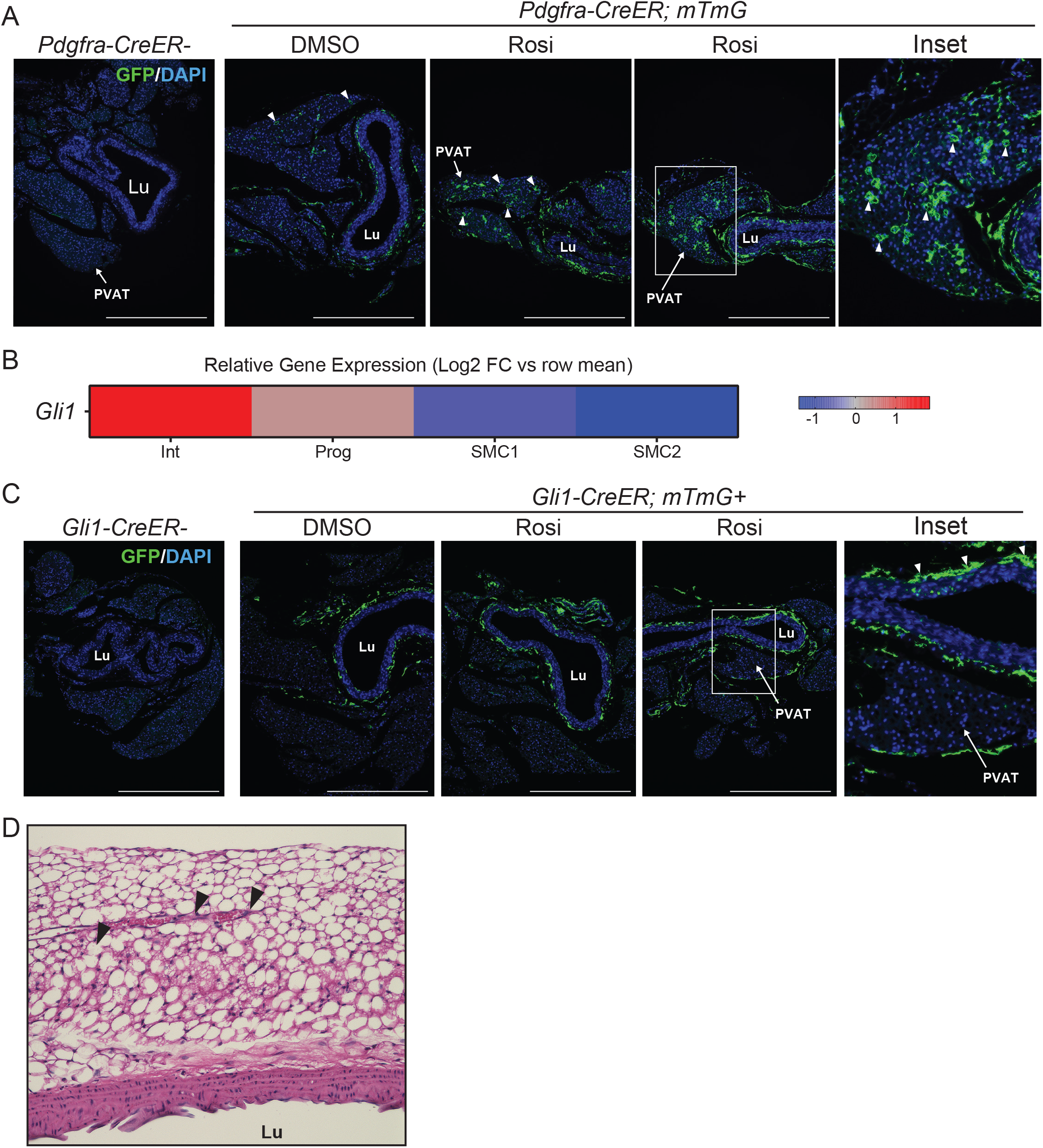
Analysis of aortic PVAT maintenance in adult mice (Related to Fig. 7) **(A)** Immunostaining of GFP (green) and DAPI (blue) in sections of thoracic aorta from *Pdgfra-CreER*−; mTmG+ (control) and *Pdgfra-CreER*+; mTmG+ mice following a 5 day pulse of Tamoxifen and a 2.5 week treatment of DMSO or Rosiglitazone. Arrowheads show GFP-labeled adipocytes (n=1 Cre−; n=4 DMSO; n=5 Rosi; scale bar, 543.5um). **(B)** Heat map showing *Gli1* expression levels in sorted cell populations. **(C)** Immunostaining of GFP (green) and DAPI (blue) in sections of thoracic aorta from *Gli1-CreER−*; mTmG+ and *Gli1-CreER*+; mTmG+ mice following a 5-day pulse with Tamoxifen and 2.5 week treatment with DMSO or Rosiglitazone. Arrowheads indicate GFP labeled cells (n=1 Cre−; n=3 DMSO; n=3 Rosi scale bar, 543.5um). **(D)** H&E staining of a longitudinal section of adult thoracic aorta. Arrowhead shows adventitial blood vessel (scale bar, 362.3um). Lu: vessel lumen.

## Notes

### Competing Interest Statement

The authors have declared no competing interest.

